# CaRPOOL: A Pooled Calcium-Recording CRISPR Screening Platform Identifies CCR7 as a Modulator of Cellular Osmomechanosensing

**DOI:** 10.1101/2025.11.27.690909

**Authors:** Miao Ouyang, Jianhui Wang, Xiaoguang Luo, Ruilin Tian

## Abstract

Cells must continuously sense and respond to environmental changes by translating physical and chemical cues into intracellular signals. However, systematic discovery of genes governing these sensory processes has been limited by the transient nature of signaling events and the low throughput of measurement assays. Here, we present CaRPOOL, a pooled, high-throughput genetic screening platform that integrates the calcium-activity recorder CaMPARI2 with CRISPR interference (CRISPRi), enabling stable capture of transient calcium signals for genome-scale functional screening. Using osmomechanical stimulation as a model, we demonstrate that CaMPARI2 photoconversion faithfully reports stimulus-dependent calcium responses and supports pooled fluorescence-activated cell sorting (FACS)–based screening. A CRISPRi library targeting membrane-associated genes identified both known and previously uncharacterized regulators of mechanotransduction, including the chemokine receptor CCR7. Mechanistic analyses revealed that CCR7 promotes osmomechanical calcium signaling through a PIEZO1-dependent Gαs–cAMP–PKA pathway, establishing it as a mechanically responsive GPCR. Notably, osmotic stress upregulated *CCR7* expression in immune cell lines and enhanced mechanical responsiveness, suggesting a role in immunomechanical adaptation. Together, these findings introduce a broadly applicable platform for high-throughput discovery of genes controlling dynamic signaling responses and reveal a GPCR–ion channel crosstalk mechanism in mechanotransduction with potential implications for immune cell mechanoadaptation.

## INTRODUCTION

Cells continuously sense and respond to environmental stimuli, translating physical and chemical cues into intracellular signals that maintain homeostasis and guide development and adaptation [1–3]. These stimuli—including mechanical forces[4, 5], temperature[6, 7], light [8], and chemical ligands[9, 10]—are primarily detected by membrane proteins such as G protein–coupled receptors (GPCRs) and ion channels that act as molecular sensors coupling external inputs to intracellular signaling cascades[11]. The identification of these sensory molecules, along with their regulatory components that modulate receptor activity, localization, and downstream signaling, has been central to understanding how organisms perceive and adapt to their surroundings.

Genetic screening has been instrumental in uncovering key genes mediating cellular responses to environmental stimuli. A landmark example is the discovery of PIEZO1 as a mechanosensitive ion channel, achieved through systematic siRNA knockdown in mouse neuroblastoma cells followed by whole-cell patch-clamp recordings to measure mechanically activated currents in response to cell poking—a laborious process requiring manual electrophysiological analysis of hundreds to thousands of individually transfected cells[12]. Similarly, identification of TRPV1 as the capsaicin and heat receptor relied on expression cloning in HEK293 cells combined with calcium imaging of individual clones, an approach that required screening hundreds to thousands of cDNA pools one by one[6]. More recently, efforts have sought to scale sensory screens using automation and genetic arrays. A screen for cold-sensing receptors in *Caenorhabditis elegans* employed a custom real-time PCR thermocycler to deliver controlled temperature stimuli while monitoring calcium responses across arrayed mutants, ultimately identifying GLR-3 as a cold sensor[13]. Likewise, Xu et al. engineered a microfluidic platform that applied shear stress stimulation in 384-well format to identify the flow-responsive GPCR GPR68[14]. While these pioneering examples illustrate the power of genetic screening in sensory biology, they also underscore a persistent limitation: most rely on low-throughput readouts or specialized instrumentation that hinder large-scale, unbiased discovery.

The fundamental challenge lies in capturing transient signaling events—which typically last seconds to minutes—in a format compatible with pooled genetic screening, where millions of cells carrying different genetic perturbations must be analyzed simultaneously. Traditional real-time sensors require continuous observation during stimulation, making them incompatible with pooled screening workflows. Endpoint assays such as transcriptional reporters can work in pooled formats but miss the immediate, dynamic signaling events that define cellular sensing. Calcium signaling, which serves as a nearly universal readout of cellular activation across diverse stimuli, would be an ideal screening parameter if its transient dynamics could be stably recorded.

CRISPR-based genetic screening has emerged as a powerful high-throughput approach to systematically dissect gene function and identify genetic determinants of complex phenotypes[15–18]. Among these, CRISPR interference (CRISPRi) leverages a catalytically dead Cas9 (dCas9) fused to a transcriptional repressor to silence gene expression[19, 20]. This is achieved when dCas9 is guided by sgRNAs targeting regions proximal to the transcription start site (TSS) of target genes. A widely adopted CRISPRi strategy employs dCas9 fused to the Krüppel-associated box (KRAB) domain (dCas9-KRAB), which induces heterochromatin formation and transcriptional repression at specific genomic loci.

In this study, we developed CaRPOOL (<u>Ca</u>lcium-<u>R</u>ecording <u>Pool</u>ed screening platform), a high-throughput genetic screening platform that integrates the photoconvertible calcium-activity recorder CaMPARI2 with CRISPR screening, enabling systematic discovery of genes that govern cellular responses to external stimuli. Applying this platform to osmomechanical stimulation (hypotonic swelling-induced mechanical stress) uncovered the chemokine receptor CCR7 as a previously unrecognized regulator of mechanotransduction that operates in a PIEZO1-dependent manner through the Gαs–cAMP–PKA signaling axis. By providing a broadly applicable approach for uncovering regulators of mechanosensation and other stimulus-responsive processes, this technology expands the toolkit for elucidating how cells perceive and translate environmental forces into biological function.

## RESULTS

### CaMPARI2 Enables High-Throughput Measurement of Calcium Responses to Osmotic Mechanical Stimulus

Various extracellular stimuli, such as changes in temperature, mechanical force, osmolarity, and pH, induce cellular responses that converge in elevated cytosolic calcium concentration. Therefore, in cellular contexts where calcium flux is a primary downstream event, calcium signaling can serve as a readout for responses to external stimuli. Calcium imaging using small-molecule probes such as Fluo-4 and Fura-2, or genetically encoded calcium sensors like GCaMP[21], is a widely used approach for real-time monitoring of intracellular Ca²⁺ dynamics. However, because calcium signals are transient and dynamic, these methods rely on fluorescence microscopy or plate readers, limiting their throughput to low- or medium-scale arrayed screens and making them unsuitable for pooled, high-throughput applications.

By contrast, calcium integrators provide a means to record transient calcium activity into a stable fluorescence readout. The Ca²⁺-Activated Photoactivatable Ratiometric Integrator (CaMPARI) is a photoconvertible fluorescent protein that irreversibly switches its state from green to red fluorescence under violet light, but only in the presence of elevated Ca²⁺ [22, 23]. This property allows the recording of cellular calcium activity during a defined illumination window into a stable, integrated fluorescence parameter—the red-to-green (R/G) ratio—which can be measured at single-cell resolution by flow cytometry, thus enabling pooled high-throughput screening. The higher the CaMPARI R/G ratio, the greater the accumulated Ca²⁺ during the illumination period, reflecting stronger cellular responses to a given stimulus (Figure 1A).

**Figure 1.**
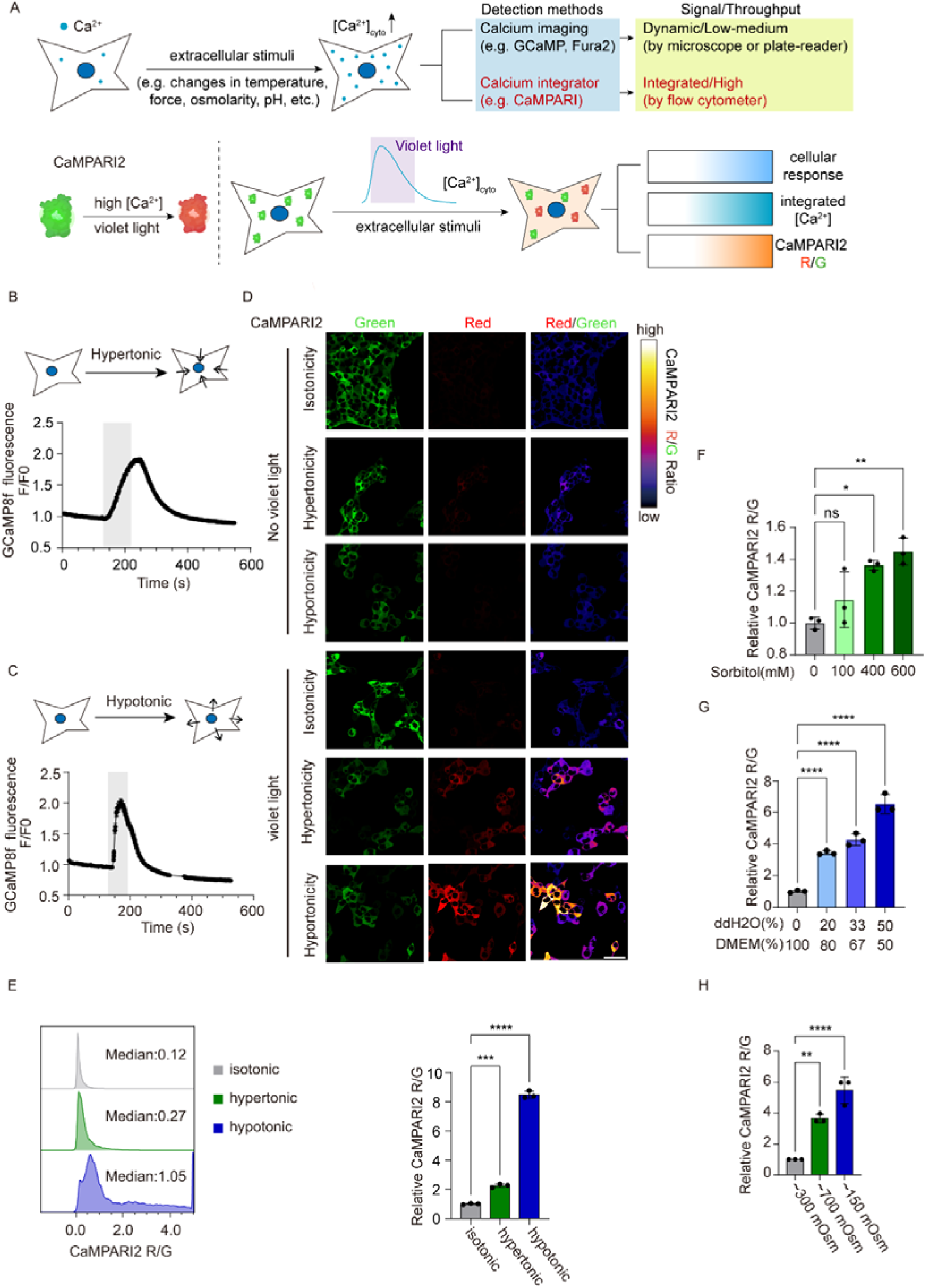
CaMPARI2 enables high-throughput measurement of calcium responses to osmotic mechanical stimulation. (A) Schematic overview of calcium detection methods. Top: Dynamic sensors (e.g., GCaMP, Fura-2) enable real-time monitoring but are limited to low- or medium-throughput applications. Calcium integrators (e.g., CaMPARI) convert transient calcium signals into stable fluorescence outputs measurable by flow cytometry for high-throughput screening. Bottom: CaMPARI2 irreversibly switches from green to red fluorescence upon violet illumination in the presence of elevated [Ca²⁺], with the red-to-green (R/G) ratio reflecting integrated calcium activity. (B–C) Real-time calcium responses of HEK293T cells to (B) hypertonic (400 mM sorbitol in DMEM,∼700mOSM) or (C) hypotonic (1:1 DMEM:ddH₂O,∼150mOSM) stimulation monitored by GCaMP8f imaging. Gray shading indicates stimulation period. Traces show mean ΔF/F₀ ± SEM. n = 79 cells (B), n = 126 cells (C). (D) CaMPARI2 photoconversion in response to osmotic stimuli. Cells were exposed to isotonic (DMEM), hypotonic (1:1 DMEM:ddH₂O,∼150mOsm), or hypertonic (DMEM + 400 mM sorbitol,∼700mOsm) conditions during 3 min violet illumination (405 nm) or kept in dark (No violet). Representative images show green (unconverted, 488nm exitation), red (photoconverted, 561nm exitation), and red/green ratio. Scale bar, 40 μm. (E) Flow cytometry quantification of CaMPARI2 R/G ratios. Left: Representative histograms. Right: Relative R/G ratios normalized to isotonic control. Data are mean ± SD (n = 3 biological replicates). ***P < 0.001, ****P < 0.0001. One-way ANOVA. (F-G) Osmolarity-dependent CaMPARI2 responses measured by flow cytometry. Cells were exposed to indicated hypotonic (F) or hypertonic (G) treatments. Data are mean ± SD (n = 3 biological replicates). ns, not significant,*P < 0.05, **P < 0.01. One-way ANOVA. (H) CaMPARI2 responses to defined osmotic solutions (sorbitol + water + CaCl₂). Cells were treated with hypertonic (∼700 mOsm), isotonic (∼300 mOsm), hypotonic (∼150 mOsm), or ddH₂O during violet illumination. Relative R/G ratios normalized to ddH₂O are shown as mean ± SD (n = 3 biological replicates). **p < 0.01, ****p < 0.0001. One-way ANOVA.

We sought to determine whether CaMPARI could be applied to high-throughput screening of cellular responses to mechanical stimulation. To deliver uniform mechanical stimulation to large populations of cells in a scalable manner, we employed osmotic mechanical stress. By altering extracellular osmolarity, cells undergo volume changes and membrane deformation, generating mechanical force without the need for complex physical devices. This approach is commonly used to mimic mechanical stress in cellular systems.

HEK293T cells were selected for these experiments due to their robust growth, ease of handling, and their well-established utility as a model system for molecular and cellular studies. We first verified whether HEK293T cells generate calcium responses to osmotic mechanical stimulation. Cells were treated with a hypertonic solution containing sorbitol, a non-ionic osmolyte, or with a hypotonic solution prepared by diluting the culture medium with water. Using the calcium indicator GCaMP8f, we detected robust and osmolarity-dependent increases in cytosolic calcium concentration in HEK293T cells by both fluorescence imaging (Figures 1B-C) and microplate reader (Figures S1A-B), consistent with previous reports[24–28].

We then tested whether CaMPARI could report cellular responses to osmomechanical stimulation. To this end, we generated a HEK293T cell line stably expressing CaMPARI2, an improved version of CaMPARI that exhibits brighter fluorescence, higher green-to-red photoconversion contrast, faster calcium unbinding kinetics, and reduced dark photoconversion[22, 23]. Cells were subjected to hypertonic or hypotonic treatments while being simultaneously exposed to violet light for 3 minutes. Both fluorescence microscopy and flow cytometry revealed significant increases in the CaMPARI2 red-to-green (R/G) ratios under hypertonic and hypotonic conditions compared to isotonic or no–violet light controls, with responses scaling in an osmolarity-dependent manner (Figures 1D–G). Notably, hypotonic stimulation elicited a stronger calcium response than hypertonic stress. Based on this observation, we selected a 1:1 H₂O-to-medium mixture as the osmomechanical stimulation condition for downstream experiments.

To exclude potential ionic effects, we prepared defined solutions composed only of sterile water, sucrose, and CaCl₂, with theoretical osmolarities approximating hypertonic (∼700 mOsm/L), isotonic (∼300 mOsm/L), and hypotonic (∼150 mOsm/L) conditions. CaCl₂ was included at a constant concentration (2.5 mM) across all three conditions to maintain extracellular calcium availability for channel-mediated influx, while sucrose served as the sole osmolyte. Comparable CaMPARI2 responses were observed under these conditions, confirming that the calcium increases were primarily driven by osmotic pressure rather than ionic composition (Figure 1H).

In summary, these results demonstrate that HEK293T cells exhibit robust calcium response to both hypertonic and hypotonic stimulation and establish CaMPARI2 as a sensitive and high-throughput tool for measuring cellular responses to osmomechanical stimuli.

### CaMPARI2-Based CRISPRi Screen Reveals Known and Novel Regulators of Osmomechanical Signaling

We next integrated CaMPARI2 with high-throughput pooled CRISPR interference (CRISPRi) screening—termed CaRPOOL (Calcium-Recording Pooled screening platform)—to systematically identify genes involved in osmomechanical signaling. To this end, we established a HEK293T cell line stably expressing both the CRISPRi machinery (dCas9–KRAB) [29]and CaMPARI2 and transduced it with a lentiviral sgRNA library targeting 2,418 membrane-associated genes (H6 library from CRISPRi-v2) [30]along with 250 non-targeting controls (Methods). We focused on membrane proteins because known mechanotransducers, such as ion channels and GPCRs, are predominantly membrane-localized.

Following selection and expansion, cells were exposed to hypotonic stimulation and 3 minutes of violet light to induce CaMPARI2 photoconversion. A portion of the population was retained as an input control, while the remainder was sorted by fluorescence-activated cell sorting (FACS) based on the CaMPARI2 red/green (R/G) ratio. For each sample, cells corresponding to at least 1000-fold over the library coverage were sorted per replicate. Cells with low R/G ratios—indicative of reduced calcium responses—were isolated by FACS (bottom 15%). The representation of each sgRNA was quantified in both input and sorted populations by next-generation sequencing (NGS). Comparison of sgRNA abundance between populations enabled identification of genes important for osmomechanical sensing, whose sgRNAs would be enriched in the sorted low R/G population (Figure 2A, Table S2).

**Figure 2.**
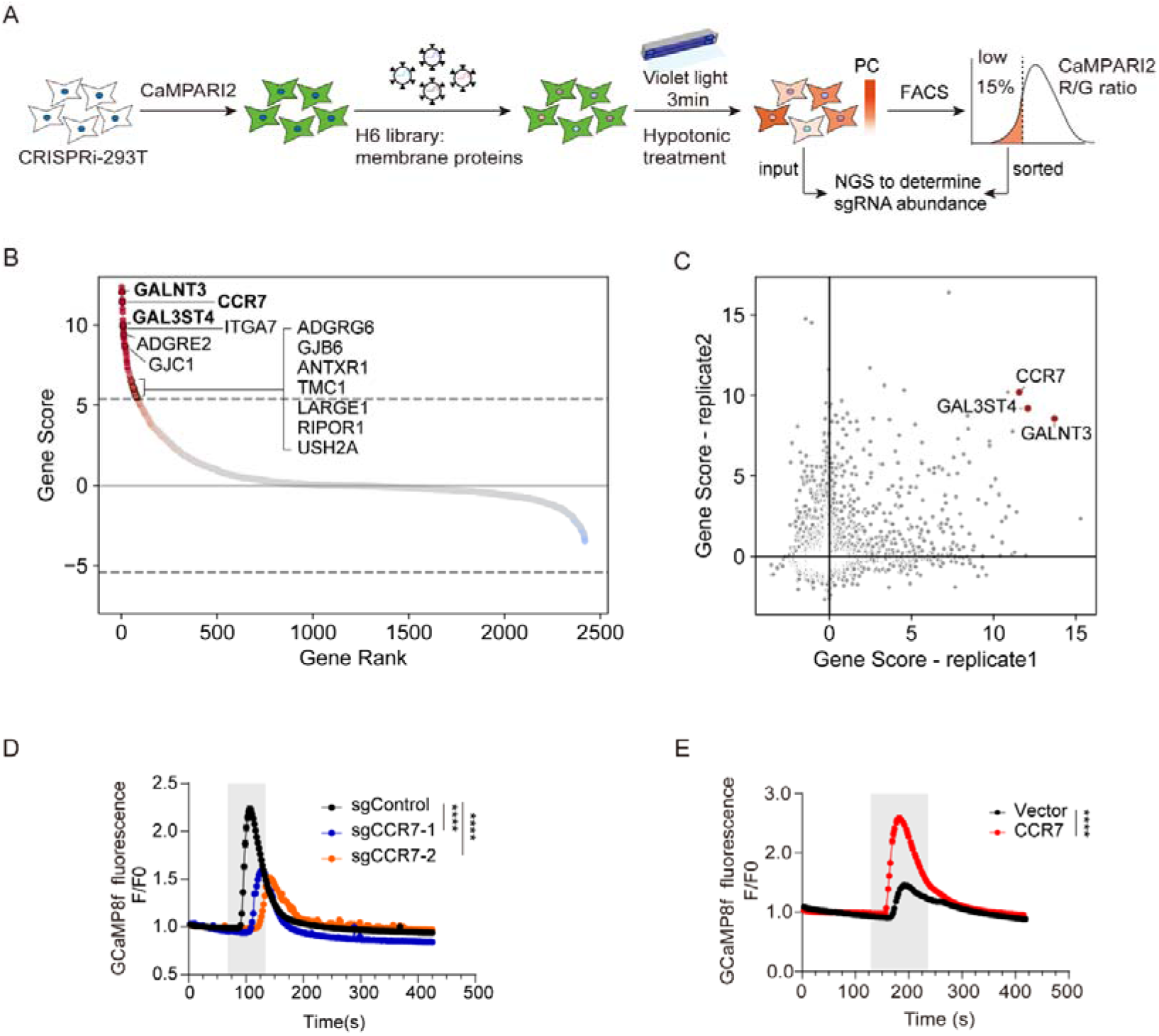
CaMPARI2-based CRISPRi screen reveals known and novel regulators of osmomechanical signaling. (A) Schematic of the CaMPARI2-based pooled CRISPRi screening workflow. HEK293T cells stably expressing CRISPRi machinery (dCas9–KRAB) and CaMPARI2 were transduced with a lentiviral sgRNA library targeting 2,418 membrane-associated genes (H6 library). Following selection and expansion, cells were subjected to hypotonic stimulation with 3 min violet light (405 nm) illumination to induce CaMPARI2 photoconversion. Cells with low CaMPARI2 R/G ratios (bottom 15%) were isolated by FACS, and sgRNA representation in input and sorted populations was quantified by next-generation sequencing (NGS) to identify genes regulating osmomechanical calcium responses. (B) Gene-level scores from the CRISPRi screen. Genes are ranked by their scores from the MAGeCK-iNC pipeline. Genes with positive scores indicate enrichment of their sgRNAs in the low R/G population, and vice versa. Top hits include known mechanotransduction genes and novel candidates. Dashed horizontal lines indicate gene score threshold for FDR < 0.1. (C) Comparison of gene scores from replicate 1 (x-axis) and replicate 2 (y-axis) shows reproducible hits, with representative candidates highlighted. (D) Validation of screen hits by calcium imaging. HEK293T cells expressing two distinct sgRNAs targeting *CCR7* or sgControl were transduced with GCaMP8f and subjected to hypotonic stimulation. Traces show mean F/F₀ ± SEM (n = 409 cells for sgControl, n = 407 cells for sgCCR7-1, n = 424 cells for sgCCR7-2). Gray shading indicates stimulation period. ****P < 0.0001. One-way ANOVA. (E) *CCR7* overexpression enhances calcium responses to hypotonic stimulation. HEK293T cells transfected with *CCR7,* or empty vector were subjected to hypotonic treatment, and GCaMP8f fluorescence was monitored. Traces show mean F/F₀ ± SEM (n = 177 cells for vector, n = 239 cells for *CCR7* overexpression). Gray shading indicates stimulation. ****P < 0.0001. Unpaired two-tailed Student’s t-test.

Our screen recovered several genes with known or plausible functions in mechanical signaling (Figure 2B), including *TMC1* and *USH2A*, which function in cochlear hair cells to mediate auditory mechanotransduction—TMC1 as a pore-forming component of the mechanosensory channel [31, 32] and USH2A as a structural element stabilizing stereocilia bundles[33]. We also identified *ITGA7* [34, 35] and *LARGE1*[36], which facilitate and maintain cell–matrix adhesion essential for force transmission; *ADGRG6*, *ADGRE2*, and *ANTXR1*, adhesion GPCRs that sense extracellular matrix stiffness or tension to activate downstream signaling[37]; *RIPOR1*, which regulates actin cytoskeleton dynamics in response to mechanical cues[38]; and *GJB6* and *GJC1*, gap junction proteins that coordinate intercellular communication during mechanically induced responses[39, 40]. These results validated the reliability of our screening approach.

We also identified genes not previously linked to osmomechanosensing and selected three top hits showing consistent phenotypes across two screen replicates for validation: *CCR7*, *GALNT3*, and *GAL3ST4*. *CCR7* encodes a chemokine receptor[41], while GALNT3 and GAL3ST4 are involved in glycosylation pathways[42, 43]. We cloned sgRNAs for each gene and individually validated their phenotypes by calcium imaging using GCaMP8f, orthogonal to our screening approach. Knockdown of each gene (Figure S2A) resulted in reduced HEK293T cell responses to hypotonic stimulation, albeit to varying degrees (Figures 2D and S2B-C), confirming the reliability of the screen.

Knockdown of *CCR7* resulted in the most significant reduction in cellular response to hypotonic stimulation (Figure 2D). Conversely, ectopic overexpression of *CCR7* markedly enhanced calcium responses to hypotonic stimulation, as measured by GCaMP8f fluorescence (Figure 2E). Notably, overexpression of *CCR6*, a closely related chemokine receptor, did not increase osmomechanical calcium signaling (Figure S2D), indicating a specific role for CCR7 in this process.

While CCR7 is canonically activated by the cytokines CCL19 and CCL21, its role in modulating osmomechanical response appears ligand-independent. Mutations (K50A and W114A) that abolish CCR7’s binding to CCL21 do not alter its ability to enhance the osmomechanical response (Figure S2E-F), suggesting a distinct activation mechanism in this context

In summary, our results highlight the CaMPARI2-based CRISPRi screen as an effective and scalable platform for discovering regulators of calcium responses to environmental stimuli and uncover *CCR7* as a novel modulator of osmomechanical signal transduction, functioning independently of its canonical ligand binding.

### CCR7 Regulates Osmomechanical Calcium Signaling Through PIEZO1-Mediated Calcium Influx

We next investigated the mechanism by which CCR7 regulates osmomechanical calcium signaling. As a GPCR, CCR7 may modulate cytosolic calcium levels through two principal pathways: (1) the Gαq–PLCβ–IP₃/DAG signaling cascade, in which inositol 1,4,5-trisphosphate (IP₃) binds to IP₃ receptor type 3 (ITPR3) on the endoplasmic reticulum (ER), triggering calcium release from ER stores into the cytosol[44], or (2) modulation of plasma membrane ion channels via G-protein coupling, leading to extracellular calcium influx[45, 46].

To determine whether hypotonic stimulation mobilizes ER calcium stores, we knocked down *ITPR3*, which encodes the IP₃ receptor responsible for ER calcium release (Figure S3A). GCaMP8f calcium imaging revealed that *ITPR3* knockdown did not affect HEK293T cell responses to hypotonic stimulation (Figure 3A), indicating that ER calcium stores do not contribute significantly to the osmomechanical response. In contrast, removal of extracellular calcium using calcium-free buffer dramatically reduced the hypotonic-induced calcium response (Figure 3B). These results demonstrate that the calcium increase induced by hypotonic mechanical stimulation in HEK293T cells primarily results from extracellular calcium influx.

**Figure 3.**
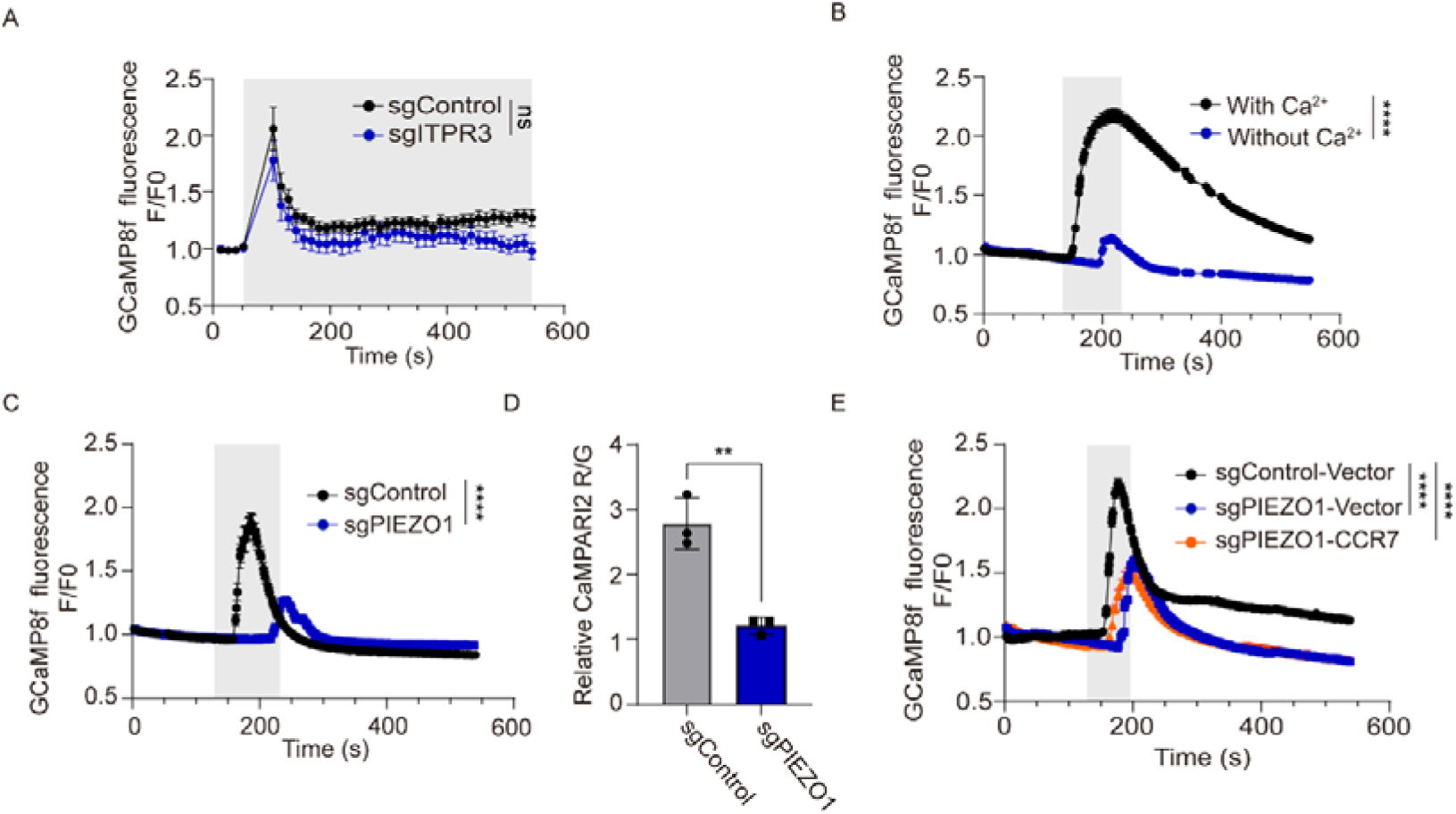
CCR7 modulates the response to hypoosmotic stimulation via PIEZO1. (A) Knockdown of IP₃ receptor 3 (*ITPR3*) does not affect the calcium response to hypotonic stimulation. GCaMP8f fluorescence traces of HEK293T cells expressing sgControl or sgITPR3 show similar calcium transients upon hypotonic treatment, measured by microplate reader. Traces represent mean F/F₀ ± SEM (n = 8 biological replicates). ns, not significant. Unpaired two-tailed Student’s t-test. (B) Hypotonicity-induced calcium influx depends on extracellular Ca²⁺. GCaMP8f fluorescence responses of HEK293T cells in isotonic buffer containing Ca²⁺ (130 mM NaCl + 2.5 mM CaCl₂) or in calcium-free buffer (no CaCl₂), measured by fluorescence imaging. Traces show mean ΔF/F₀ ± SEM (n = 219 cells for with Ca²⁺ condition, n = 259 cells for without Ca²⁺ condition). Gray shading indicates stimulation period. ****P < 0.0001. Unpaired two-tailed Student’s t-test. (C) *PIEZO1* knockdown abolishes hypotonicity-induced calcium responses. GCaMP8f fluorescence responses of HEK293T cells expressing sgControl or sgPIEZO1 exposed to hypotonic medium (1:1 DMEM:ddH₂O, ∼150mOsm), measured by fluorescence imaging. Traces show mean F/F₀ ± SEM (n = 94 cells for sgControl, n = 299 cells for sgPIEZO1). ****P < 0.0001. Unpaired two-tailed Student’s t-test. (D) CaMPARI2 assay confirms reduced calcium response in *PIEZO1*-depleted cells. Relative CaMPARI2 R/G ratios of sgPIEZO1 and sgControl cells after 3 min violet illumination during hypotonic stimulation. Data are mean ± SD (n = 3 biological replicates). **P < 0.01. Unpaired two-tailed Student’s t-test. (E) *CCR7* overexpression fails to restore calcium influx in *PIEZO1*-knockdown cells. GCaMP8f traces of sgPIEZO1 cells transfected with vector or *CCR7* are shown alongside sgControl-Vector cells, measured by fluorescence imaging. Traces represent mean F/F₀ ± SEM (n = 312 cells for sgControl-Vector, n = 240 cells for sgPIEZO1-Vector, n = 196 cells for sgPIEZO1-CCR7). Gray shading indicates stimulation. ****P < 0.0001. One-way ANOVA.

PIEZO1 is a mechanosensitive cation channel responsible for Ca²⁺ influx in diverse mechanical responses across multiple cell types[47–51]. We therefore examined whether PIEZO1 mediates calcium influx under osmomechanical stimulation. Knockdown of *PIEZO1* in HEK293T cells significantly reduced the hypotonic calcium response, as determined by both GCaMP8f imaging and CaMPARI2 assays (Figures 3C-D and S3B), indicating a major role for PIEZO1 in mediating the HEK293T hypotonic response.

We next investigated whether CCR7 influences the hypotonic response through PIEZO1. We overexpressed *CCR7* in *PIEZO1*-knockdown HEK293T cells and performed GCaMP8f calcium imaging. In *PIEZO1*-knockdown cells, *CCR7* overexpression no longer enhanced the hypotonic response (Figure 3E), suggesting that CCR7’s effect is PIEZO1-dependent. To exclude potential residual PIEZO1 activity in knockdown cells, we generated a *PIEZO1* knockout HEK293T cell line (Figure S3C). These cells exhibited near-complete loss of calcium response to hypotonic treatment (Figure S3D). Consistently, CCR7 overexpression failed to enhance the hypotonic response in *PIEZO1*-KO cells (Figure S3D), confirming that CCR7 modulates the HEK293T hypotonic response in a PIEZO1-dependent manner.

Together, these findings indicate that CCR7 regulates cellular responses to osmomechanical stimuli through PIEZO1-mediated extracellular calcium influx.

### CCR7 Modulates Osmomechanical Calcium Signaling via the Gαs-cAMP-PKA Pathway

A recent study reported that protein kinase A (PKA) directly phosphorylates PIEZO1 to enhance its mechanosensitivity[52, 53]. Because PKA can be activated downstream of GPCR signaling through the Gαs–cAMP pathway, we examined whether CCR7 modulates PIEZO1 signaling via this mechanism.

Knockdown of Gαs (encoded by *GNAS*) using CRISPRi in HEK293T cells markedly reduced the calcium response to hypotonic stimulation and almost completely abolished the enhancement induced by CCR7 overexpression (Figures 4A and S4A), indicating that CCR7 acts through Gαs-dependent signaling. A CCR7 mutant carrying substitutions (N255W, F256R, R258E, R154W) in the predicted CCR7–G protein interaction surface within ICL2 and ICL3 lost its ability to enhance the osmomechanical response (Figures S4B-C), further supporting a direct CCR7–G protein interaction in osmomechanical regulation.

Because Gαs activates adenylate cyclase (AC) to produce cyclic adenosine monophosphate (cAMP), we next tested whether hypotonic stimulation elevates intracellular cAMP levels in HEK293T cells. Using the GloSensor™ cAMP assay, we observed a sustained increase in cAMP following hypotonic stimulation, whereas isotonic treatment showed little change (Figure 4B). Forskolin, an AC activator, served as a positive control (Figure S4D). Consistent results were obtained using the genetically encoded cAMP sensor GFlamp2[54, 55], which showed osmolarity-dependent increases in cAMP (Figure S4E), confirming AC activation.

**Figure 4.**
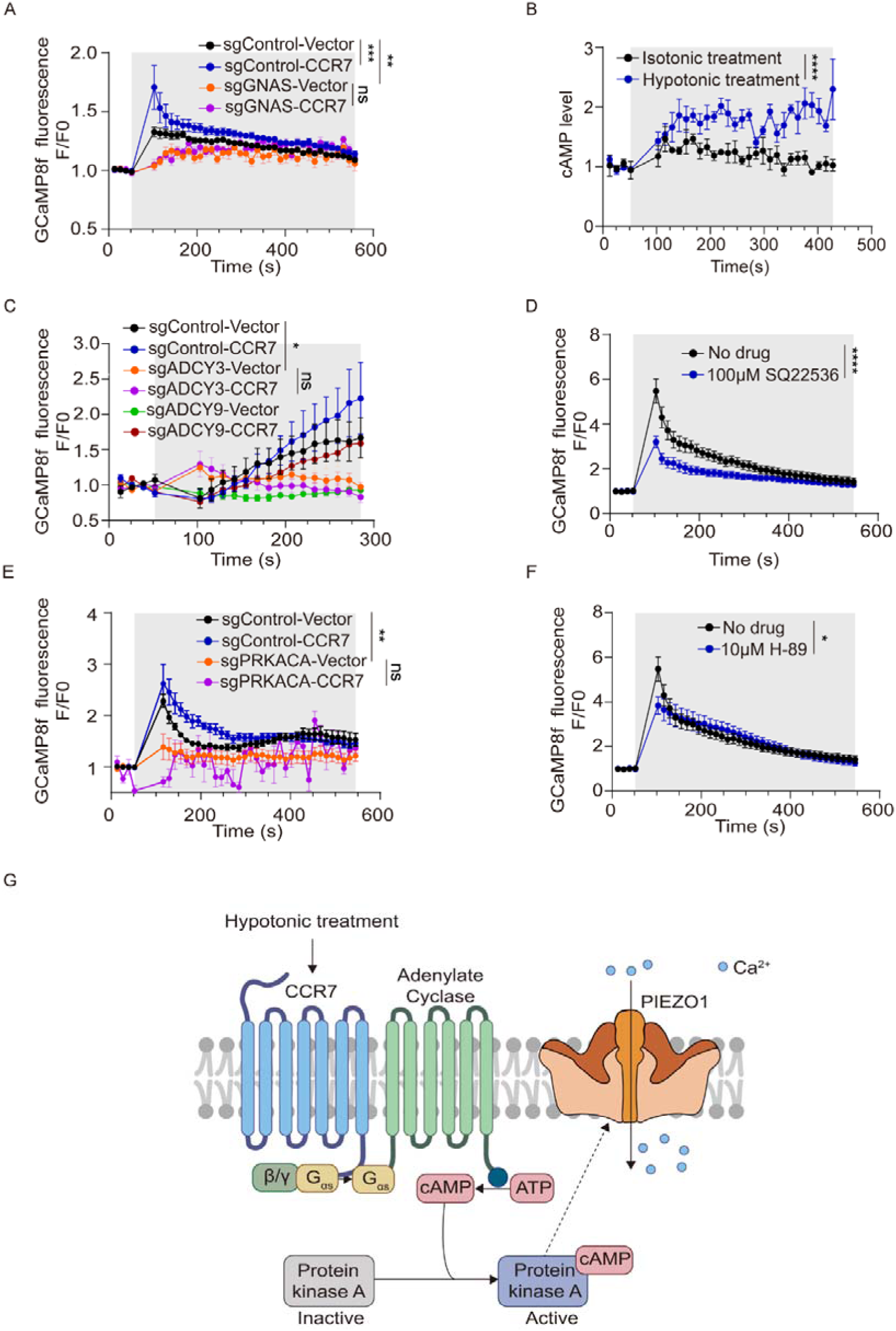
CCR7 modulates PIEZO1-mediated calcium influx through the Gαs -cAMP-PKA pathway. (A) CCR7-mediated calcium response modulation requires Gαs signaling. HEK293T cells expressing sgControl or sgGNAS with vector or *CCR7* were subjected to hypotonic stimulation, measured by microplate reader. Traces show mean F/F₀ ± SEM (n = 10 biological replicates). Gray shading indicates stimulation. ns, not significant, **P < 0.01, ***P < 0.001. One-way ANOVA. (B) Hypotonic stimulation increases intracellular cAMP levels. Cells expressing the cAMP biosensor pGloSensor-22F were monitored during isotonic or hypotonic treatment by microplate reader. Traces show mean cAMP levels ± SEM (n = 3 biological replicates). Gray shading indicates stimulation. ****P < 0.0001. Mann-Whitney U test. (C) Adenylyl cyclase (*ADCY3*) knockdown abolishes CCR7-mediated enhancement of calcium responses. GCaMP8f traces of cells expressing sgControl or sgADCY3 with vector or CCR7, measured by microplate reader. Traces show mean F/F₀ ± SEM (n = 5 biological replicates). Gray shading indicates stimulation. ns, not significant, *P < 0.05. One-way ANOVA. (D) Adenylyl cyclase inhibitor SQ22536 blocks hypotonicity-induced calcium influx. HEK293T cells pretreated with 100 μM SQ22536 or vehicle were subjected to hypotonic stimulation, measured by microplate reader. Traces show mean F/F₀ ± SEM (n = 9 biological replicates). Gray shading indicates stimulation. ****P < 0.0001. Mann-Whitney U test. (E) Protein kinase A catalytic subunit (*PRKACA*) knockdown reduces calcium responses to hypotonic stimulation and blocks CCR7-mediated enhancement of calcium responses. GCaMP8f traces of cells expressing sgControl or sgPRKACA with vector or *CCR7*, measured by microplate reader. Traces show mean F/F₀ ± SEM (n = 4 biological replicates). Gray shading indicates stimulation. ns, not significant, **P < 0.01. One-way ANOVA. (F) PKA inhibitor H-89 suppresses hypotonicity-induced calcium influx. HEK293T cells pretreated with 10 μM H-89 or vehicle were subjected to hypotonic treatment, measured by fluorescence imaging. Traces show mean F/F₀ ± SEM (n = 9 biological replicates). Gray shading indicates stimulation. *P < 0.05. Mann-Whitney U test. (G) Proposed model: Osmomechanical stimulation activates CCR7-Gαs-AC signaling, producing cAMP that activates PKA to phosphorylate and enhance PIEZO1 activity, leading to increased Ca²⁺ influx.

Furthermore, knocking down *ADCY3*, a major AC isoform expressed in HEK293T cells, substantially reduced the hypotonic calcium response and prevented CCR7 from enhancing it (Figures 4C and S4F). Similarly, inhibiting AC activity with the small-molecule inhibitor SQ22536 significantly diminished the calcium response (Figure 4D).

Since cAMP activates PKA, we next asked whether PKA mediates CCR7’s effect. Inhibition of PKA, either by silencing *PRKACA* (encoding the PKA catalytic subunit) or by treatment with H-89, significantly reduced calcium responses to hypotonic stimulation (Figures 4E-F and S4G). *PRKACA* knockdown also eliminated the potentiating effect of CCR7 overexpression (Figure 4E). Thus, PKA activity is required for CCR7-dependent enhancement of osmomechanical calcium influx.

Together, these findings reveal that CCR7 enhances osmomechanical calcium Signaling through the Gαs–AC–cAMP–PKA signaling axis (Figure 4G).

### CCR7 Expression is Induced by Osmomechanical Stimulation and Enhances Calcium Signaling in Immune Cell Lines

CCR7 is highly expressed in immune cells, including T cells and dendritic cells (DCs), where it regulates migration and activation during immune responses[56–59]. Because these cells encounter variable mechanical environments during circulation and tissue infiltration, we asked whether they respond to osmomechanical cues and whether CCR7 participates in this process.

We first sought to determine how immune cells respond to osmomechanical stimulation at the transcriptomic level. To this end, we treated Jurkat T cells, a human T-lymphocyte cell line, with hypotonic or isotonic medium for 6 hours and performed RNA-sequencing (Figure 5A). Hypotonic stimulation induced widespread transcriptional changes, with 932 genes upregulated and 539 downregulated (Figure 5B). Gene ontology analysis revealed enrichment of pathways related to T cell activation, lymphocyte differentiation, and Wnt signaling (Figures 5C and 5D), indicating that osmomechanical stress activates transcriptional programs associated with T cell functional regulation.

**Figure 5.**
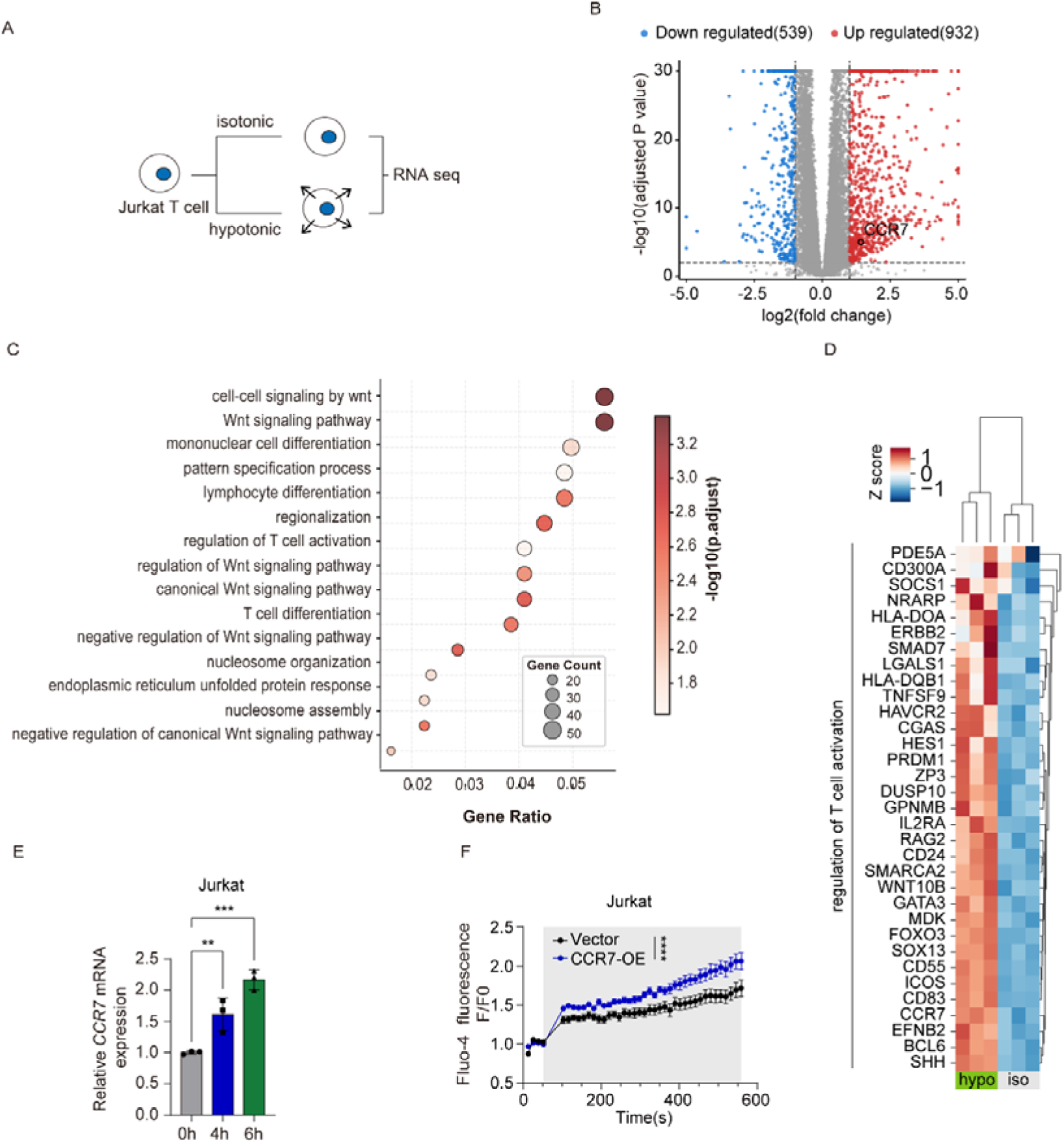
Hypotonic stimulation upregulates *CCR7* and enhances calcium responses in immune cells. (A) Schematic of RNA-seq experimental design. Jurkat T cells were exposed to isotonic or hypotonic medium for 6 hours, followed by RNA sequencing. (B) Volcano plot showing differentially expressed genes in Jurkat T cells after hypotonic treatment. Blue dots represent downregulated genes (n = 539), red dots represent upregulated genes (n = 932), and gray dots represent genes with no significant change. *CCR7* is highlighted as significantly upregulated. Thresholds: |log2(fold change) | > 1 and adjusted P value < 0.05. (C) Gene Ontology (GO) enrichment analysis of differentially expressed genes. Top 15 biological process terms ranked by adjusted P value are shown. Dot size represents gene count, and color intensity indicates −log10(adjusted P value). (D) Heatmap showing expression of genes related to regulation of T cell activation. Z-score normalized expression levels in Jurkat cells under isotonic (iso) or hypotonic (hypo) conditions are displayed. (E) Hypotonic stimulation upregulates *CCR7* mRNA expression in Jurkat cells. Relative *CCR7* mRNA levels at 0, 4, and 6 hours after hypotonic treatment, measured by qRT-PCR. Data are mean ± SD (n = 3 technical replicates). **P < 0.01, ***P < 0.001. One-way ANOVA. (F) *CCR7* overexpression enhances calcium responses to hypotonic stimulation in Jurkat T cells. Fluo-4 fluorescence traces of cells transfected with empty vector or *CCR7*, measured by microplate reader. Traces show mean F/F₀ ± SEM (n = 8 biological replicates). Gray shading indicates stimulation. ****P < 0.0001. Unpaired two-tailed Student’s t-test.

Among the upregulated genes, *CCR7* expression was significantly increased under hypotonic conditions (Figure 5B). This increase was corroborated by quantitative PCR, which confirmed higher CCR7 mRNA levels (Figure 5E), and by flow cytometry following immunostaining, which demonstrated elevated CCR7 protein levels on the surface of Jurkat cells (Figures S5A-B) after hypotonic treatment, consistent with previous observations that mechanical confinement upregulates *CCR7* in immune cells[56]. These findings suggest that osmomechanical cues induce *CCR7* expression as part of a mechanoadaptive transcriptional response.

We next examined whether increased *CCR7* expression enhances the cellular response to hypotonic stress. Overexpression of CCR7 significantly augmented calcium influx upon hypotonic stimulation in Jurkat T cells, as measured by Fluo-4 calcium imaging (Figure 5F). Similar phenotypes were observed in DC2.4 cells, an immortalized murine dendritic cell line used as another model system: hypotonic stimulation increased *CCR7* expression in these cells (Figure S5C), and *CCR7* overexpression further enhanced calcium influx in response to hypotonic stimulation (Figure S5D).

Together, these results show that osmomechanical stimuli upregulate *CCR7* expression in immune cells, and elevated CCR7 levels amplify calcium responses to hypotonic stress. This suggests the existence of a positive feedback mechanism in which osmomechanical activation enhances *CCR7* expression, which in turn sensitizes cells to subsequent mechanical cues, thereby facilitating rapid immunomechanical adaptation. Further validation in primary immune cells would be essential to confirm these findings in a physiological context.

## Discussion

This study establishes CaRPOOL, a FACS-based pooled screening system as a scalable platform for dissecting cellular responses to environmental stimuli. By integrating a calcium-sensitive photoconvertible reporter CaMPARI2 with CRISPR interference and flow cytometry, we we leveraged CaMPARI2’s ability to permanently record transient calcium activity as a stable fluorescence ratio, enabling its integration into a pooled CRISPRi screening workflow scalable to large cell populations. This approach overcomes a key limitation of traditional imaging-based calcium assays, enabling pooled, high-throughput genetic analysis of dynamic signaling events.

We applied this technology to osmotic mechanical stimulation as a proof-of-concept. Changes in extracellular osmolarity deform cell membranes and generate mechanical stress in a controllable manner. HEK293T cells displayed stimulus-dependent calcium responses to both hypertonic and hypotonic conditions, which CaMPARI2 captured as quantifiable fluorescence conversions. These results validated the assay’s sensitivity and demonstrated osmotic stress as an effective model for large-scale screening of mechanotransduction.

Using a CRISPRi library targeting membrane-associated genes, the system identified several known mechanosensory components and uncovered CCR7 as a previously unrecognized regulator of osmomechanical calcium signaling. Mechanistic analyses showed that upon hypotonic activation, CCR7 engages Gαs to activate adenylate cyclase and increase cAMP, leading to PKA activation, which then enhances PIEZO1 channel activity likely through direct phosphorylation. While PIEZO1 phosphorylation by PKA has been previously reported[52], further investigation is warranted to confirm this specific phosphorylation event and to identify the precise sites on PIEZO1 involved in this context. Additionally, the exact mechanism by which CCR7 itself is activated by osmomechanical stimuli requires further biochemical and structural elucidation. Future work is also needed to determine whether and how PIEZO1 might regulate CCR7.

While the screen and validation experiments in this study primarily focused on hypotonic stimulation, an important question is whether the CCR7–Gαs–PKA–PIEZO1 axis is specifically triggered by osmotic membrane deformation or whether it could also be activated by ionic strength reduction caused by the hypotonic solution. Future experiments using isosmotic ionic strength reduction (e.g., replacing NaCl with membrane-impermeant sucrose at constant osmolality) versus membrane stretch without ionic dilution (e.g., patch-clamp with negative pressure) would help delineate which physical stimulus is the relevant trigger for CCR7-dependent signaling. Such experiments would also clarify whether the CCR7–Gαs–PKA pathway represents a general mechanoadaptive mechanism applicable to diverse osmomechanical stimuli, or one specifically tuned to the membrane deformation component of hypotonic stress.

Although ion channels have been traditionally viewed as primary mechanosensors, accumulating evidence indicates that GPCRs are also involved in mediating cellular responses to physical forces. Examples include GPR68[14], H1R[48], AT1R[49–50], CysLT1R[51], GABAB[60], and several adhesion GPCRs[61–63]. CCR7 adds to this repertoire. Like many of the GPCRs listed above, CCR7 can regulate osmomechanical responses independent of its canonical ligands: mutations that disrupt binding to the canonical ligand CCL21 do not impair its ability to enhance the osmomechanical response.

The mechanically responsive properties of CCR7 are particularly relevant in the immune system, where migrating cells continually encounter shear stress, compression, and osmotic fluctuations[64, 65]. Such forces influence receptor signaling, activation thresholds, and effector functions. CCR7 is essential for lymphocyte homing and dendritic cell trafficking[66, 67]—processes that inherently involve mechanical challenge—and has been linked to mechanical modulation in both physiological and pathological contexts. Notably, we observed that osmomechanical stress upregulated *CCR7* expression in Jurkat T and dendritic DC2.4 cells, suggesting that CCR7 participates in a mechanoadaptive program where mechanical activation induces its own expression, thereby sensitizing cells to further mechanical cues. Such self-reinforcing mechanosensory circuits have precedent in other systems. For example, in cardiac fibroblasts, Piezo1 activation promotes integrin β1 clustering and cytoskeletal tension that further activates Piezo1[68] and in nucleus pulposus cells, Piezo1-driven NF-κB signaling stiffens the extracellular matrix via periostin upregulation, amplifying Piezo1 activity in return[69].

Beyond osmomechanical stimulation, the CaMPARI platform offers a broadly applicable framework for functional genomic screening of calcium-based cellular phenotypes. Recent studies have reported CaMPARI-based screening for identifying regulators of calcium activity in T cells[70, 71] and dissecting genetic modifiers of neuronal activity in human induced pluripotent stem cells (hiPSCs)-derived neurons[72].We anticipate that this approach will enable systematic, large-scale discovery of the cellular regulators governing calcium dynamics across a wide array of cell types and physiological contexts.

## MATERIALS AND METHODS

### Cell culture

All cells were maintained at 37 °C in a humidified incubator with 5% CO₂. HEK293T cells (ATCC) were cultured in Dulbecco’s Modified Eagle Medium (DMEM; Gibco, C11995500BT) supplemented with 10% fetal bovine serum (FBS; TransGen Biotech, FS301-02) and 1% penicillin–streptomycin (Aladdin, P301861). Jurkat cells were maintained in RPMI 1640 medium (Gibco, 11875093) with the same supplements. PIEZO1-knockout HEK293T cells were generously provided by Dr. Yan (Shenzhen Bay Laboratory). All cell lines were mycoplasma negative detected by MycAwayTM Plus-Color One-Step Mycoplasma Detec- tion Kit (Yeasen, 40612ES25).

### sgRNA cloning

Individual sgRNAs were cloned into the pLG15 vector using BstXI and Bpu1102I restriction sites. The pLG15 vector contains a mouse U6 promoter for sgRNA expression and an EF-1α promoter driving puromycin resistance and BFP for selection. sgRNA sequences are provided in Table S3.

### Lentivirus production for the sgRNA library

HEK293T cells were seeded in 15 cm dishes and incubated for 24 h before transfection. For each dish, 4 µg of sgRNA library plasmid and 12 µg of third-generation packaging plasmid mix (1:1:1 ratio of pMDLg/pRRE (Addgene 12251), pRSV-Rev (Addgene 12253), and pMD2.G (Addgene 12259)) were diluted in 3 mL of Opti-MEM (Gibco, 31986-07). Polyethylenimine (PEI, 2 mg/mL; Yeasen, 40816ES03) was added at 36 µL per reaction, mixed thoroughly by vortexing, and incubated for 15–20 min at room temperature before being added to the cells. Viral supernatants were harvested 48 h post-transfection, filtered through 0.45 µm filters (Millipore SLHV033RB), and used immediately or stored at −80 °C.

### Quantitative real-time PCR (qPCR)

Total RNA was extracted using the MolPure® Cell RNA Kit (Yeasen, 19231ES50). cDNA was synthesized from extracted RNA using the TransScript® One-Step gDNA Removal and cDNA Synthesis SuperMix (TransGen, AT311-03). qPCR was performed with AceQ qPCR SYBR Green Master Mix (Vazyme, CQ111-02) on an FDQ-96A fluorescence quantitative PCR system following manufacturer instructions. Gene expression was normalized to GAPDH.

### Hypotonic and hypertonic treatments

For flow-cytometry–based CaMPARI2 measurements, isotonic conditions consisted of DMEM. Hypotonic medium was prepared by diluting DMEM with an equal volume of double-distilled water (ddH₂O), while hypertonic medium was made by supplementing DMEM with 400 mM sorbitol.

For calcium imaging and microplate reader assays, the isotonic buffer contained 130 mM NaCl, 3 mM KCl, 2.5 mM CaCl₂, 0.6 mM MgCl₂, 10 mM HEPES, and 10 mM glucose, adjusted to pH 7.4 with NaOH in ddH₂O. Hypotonic buffer was prepared by diluting the isotonic buffer 1:1 (v/v) with ddH₂O, and hypertonic buffer was produced by adding 400 mM sorbitol to the isotonic buffer. The calcium-free buffer contained 130 mM NaCl, 3 mM KCl, 0.6 mM MgCl₂, 10 mM HEPES, and 10 mM glucose, adjusted to pH 7.4 with NaOH in ddH₂O, without CaCl₂.

For experiments using defined sorbitol-based osmotic solutions containing only sorbitol, water, and CaCl₂, the hypertonic solution (∼700 mOsm) consisted of 693 mM sorbitol and 2.5 mM CaCl₂ in ddH₂O; the isotonic solution (∼300 mOsm) contained 293 mM sorbitol and 2.5 mM CaCl₂ in ddH₂O; and the hypotonic solution (∼150 mOsm) included 143 mM sorbitol and 2.5 mM CaCl₂ in ddH₂O.

### Drug treatments

For adenylyl cyclase or protein kinase A inhibition, cells were seeded and incubated overnight, then treated with SQ22536 (100 µM; MCE, HY-100396) or H-89 (10 µM; MCE, HY-15979) for 2 h prior to experiments.

For adenylyl cyclase activation, cells were treated with forskolin (20 µM; SelleckChem, S2449) for 5 s immediately before stimulation.

### Measurement of the intracellular calcium concentration by fluorescence imaging

HEK293T cells were loaded with Fluo-4 AM or transfected with GCaMP8f. Prior to imaging, the culture medium was replaced with pre-warmed isotonic solution or the indicated test solutions.

Calcium fluorescence images were acquired over 10 min at room temperature using a Nikon Ti2-U fluorescence microscope (Japan) equipped with a 20× objective and a 488 nm excitation filter. In parallel, population-level fluorescence signals were measured using a microplate reader (excitation 488 nm, emission 525 nm). The experimental protocol consisted of a 2 min baseline recording in isotonic solution, followed by exposure to hypertonic or hypotonic solution for 1–2 min, and recovery in isotonic solution for an additional 6 min.

For single-cell calcium flux analysis, individual cells were outlined as regions of interest (ROI), and fluorescence intensity was quantified over time. Changes in intracellular calcium were expressed as F/F₀ = (Fₜ / F₀) × 100%, where Fₜ is fluorescence at time t and F₀ is the mean baseline fluorescence before osmotic stimulation. Statistical analyses were performed using GraphPad Prism (v9.0), calculating the mean F/F₀ from three consecutive time points around each peak calcium response.

### Measurement of the intracellular calcium concentration by microplate reader

HEK293T cells were loaded with Fluo-4 AM or transfected with GCaMP8f and seeded into black-walled 96-well plates. Each well was filled with 100 µL isotonic solution. Fluorescence measurements were performed using a microplate reader under the following conditions: excitation at 488 nm, emission at 525 nm, at room temperature.

The recording protocol consisted of an initial 1 min baseline measurement in isotonic solution, followed by addition of hypertonic or hypotonic solution and continuous recording for an additional 9 min. Relative changes in fluorescence intensity were calculated as F/F₀ = (Ft / F₀) × 100%, identical to the imaging-based method. Statistical analyses were performed in GraphPad Prism, with significance assessed by comparing mean ΔF/F₀ values derived from three time points around the peak calcium signal.

### Measurement of the intracellular calcium concentration by CaMPARI2 using flow cytometry

HEK293T cells stably expressing CaMPARI2 were seeded into 24-well plates one day prior to experiments. Before measurement, cells were incubated in pre-warmed isotonic, hypotonic, or hypertonic solution and illuminated with a 405 nm violet LED for 3 min to induce CaMPARI2 photoconversion.

Following illumination, cells were trypsinized, collected, and analyzed by flow cytometry. Fluorescence signals were recorded in the GFP (488 nm excitation) and RFP (561 nm excitation) channels, and the RFP/GFP ratio was calculated to quantify the extent of CaMPARI2 photoconversion, reflecting calcium-dependent activation. Flow cytometry data were processed using FlowJo v10.7 (Tree Star).

### CaRPOOL screen and data analysis

HEK293T cells were stably integrated with CaMPARI2 via lentiviral transduction (Plasmid sequences in Table S3). Seven days after transduction, GFP-positive cells were sorted by fluorescence-activated cell sorting (FACS; BD FACSAria SORP) to isolate CaMPARI2-expressing populations.

Lentivirus carrying the membrane-protein sgRNA library was produced as described above and transduced into CaMPARI2-CRISPRi HEK293T cells at a multiplicity of infection (MOI < 0.3). Forty-eight hours post-infection, cells were selected with puromycin (2 µg/mL) for 48 h to remove uninfected cells.

For pooled photoconversion screening, approximately 3 × 10⁷ transduced cells were plated in 150 mm dishes and illuminated with 405 nm violet light for 3 min. A fraction of 5 × 10⁶ cells was collected as the input control, while the remaining ∼3 × 10⁷ cells were subjected to FACS sorting to isolate the lowest 15% of cells based on the CaMPARI2 RFP/GFP (R/G) ratio, representing reduced calcium responses.

Genomic DNA was extracted from both input and sorted populations using DNAiso Reagent (Takara, 9770Q). sgRNA cassettes were PCR-amplified with 2× Phanta Flash Master Mix (Vazyme, P510-02) using primers oSeq_dual_p7 and oSeq_dual_p5 (sequences in Table S3) and purified using Hieff NGS DNA Selection Beads (Yeasen, 12601ES08). Libraries were sequenced on the DNBSEQ-T7 platform (MGI Tech).

Sequencing reads (FASTQ Read2) were trimmed and aligned to the reference sgRNA library using Bowtie v1.1.2 (Table S1). Guide and gene-level enrichment scores were computed using the MAGeCK-iNC pipeline [73, 74] to identify sgRNAs significantly associated with altered CaMPARI2 photoconversion phenotypes (Table S2).

### Generation of the *PIEZO1* knockout line

*PIEZO1* knockout HEK293 cells were purchased from Cyagen (STSKO240524CXB1). The knockout was generated using two sgRNAs targeting *PIEZO1* with the following protospacer sequences: sgRNA1, 5′-CCAAGGGGATGGCAGTAGCGTGG-3′; sgRNA2, 5′-AGTGAGGCCCAGAGAACGGCAGG-3′. The two sgRNAs are used cooperatively to generate a 2727 bp deletion spanning *PIEZO1* exons. Genomic DNA was extracted and used as the template for PCR amplification of the region spanning the sgRNA target sites with the primers PIEZO1-F (5′-TACCGCACACTGCAAGGTATG-3′) and PIEZO1-R (5′-GTGAGAGGACGTCCGAGAAG-3′). A sequencing primer, PIEZO1-seq (5′-CCAGAGAATGTGGCTATGCT-3′), was used for Sanger sequencing to verify the deletion.

### Measurement of cell surface levels of CCR7 by flow cytometry

Cells were stained with a fluorochrome-conjugated anti-human CCR7 antibody (BioLegend, 353203). To each sample, Cell Staining Buffer was added to bring the volume to approximately 15 mL, and the mixture was centrifuged at 350 × g for 5 minutes; the supernatant was discarded. The cells were incubated with anti-human CCR7 antibody on ice for 15–20 minutes in the dark. After incubation, samples were washed twice with at least 2 mL of Cell Staining Buffer by centrifugation at 350 × g for 5 minutes. Flow cytometry was performed on a BD FACSAria SORP, and data were analyzed using FlowJo software.

### Tango assay

HTLA cells (HEK293-derived cells stably expressing a tTA-dependent luciferase reporter and a β-arrestin2-TEV fusion protein) were maintained in DMEM supplemented with 10% fetal bovine serum (FBS) and 1% penicillin–streptomycin in a humidified incubator at 37°C with 5% CO2. Cells were plated in standard culture dishes. The following day, cells were transfected using polyethyleneimine (PEI). On day 3, transfected cells were transferred in 40 μL of medium per well into poly-L-lysine–coated, rinsed 96-well white, clear-bottom plates. On day 4, CCL21 stimulation solutions (2 ng/mL) were prepared in filter-sterilized assay buffer consisting of 20 mM HEPES and 1× HBSS at pH 7.4. On day 5, the medium and stimulant solutions were removed, and 100 μL of luciferase substrate was added per well. After incubation for 15–20 minutes at room temperature, luminescence was measured using a microplate reader.

### Measurement of intracellular cAMP

Intracellular cyclic AMP (cAMP) levels were measured using either a luminescent biosensor (pGloSensor-22F) or a fluorescent biosensor (GFlamp2).

For luminescence-based assays, HEK293T cells were transfected with the pGloSensor-22F cAMP reporter plasmid and incubated for 48 h. Cells were then subjected to isosmotic or hypoosmotic stimulation at time 0 min and immediately placed in a microplate reader at room temperature. Luminescence intensity was monitored continuously for 10 min to track dynamic cAMP changes.

For fluorescence-based assays, HEK293T cells were transfected with the GFlamp2 cAMP reporter plasmid and seeded into 96-well black-walled plates. Each well contained 100 µL isotonic solution prior to recording. Fluorescence signals (excitation 405 nm, emission 510 nm) were measured using a microplate reader under standard conditions. Cells were recorded for 1 min under isotonic conditions, followed by an additional 19 min recording after replacement with isotonic or hypotonic solution. cAMP dynamics were quantified from real-time fluorescence traces using GraphPad Prism (v9.0).

### RNA-seq and data analysis

Jurkat cells were collected after treatment, and total RNA was extracted using TRIzol Reagent (Ambion). RNA concentration and purity were measured using a Qubit 4.0 Fluorometer (Thermo Scientific), and RNA integrity was evaluated using an Agilent 2100 Bioanalyzer (Agilent Technologies). Sequencing libraries were generated using the Hieff NGS® Ultima Dual-Mode mRNA Library Prep Kit for Illumina® (Yeasen, Cat. No. 12301), following the manufacturer’s instructions. High-throughput sequencing was performed on the DNBSEQ-T7 platform (Geneplus-Shenzhen), producing at least 12 Gb of 150 bp paired-end (PE150) reads per sample.

Raw sequencing reads were aligned to the human reference genome (GRCh38) using STAR (v2.7.6a), and gene-level expression was quantified with StringTie2 (v2.0.4). Expression normalization was performed using both Fragments Per Kilobase of transcript per Million mapped reads (FPKM) and Transcripts Per Million (TPM) methods.

Differential gene expression between experimental conditions was analyzed using the DESeq2 R package v1.26.0. P-values were adjusted for multiple testing via the Benjamini–Hochberg correction method, and genes with adjusted P < 0.05 and |log₂ (fold change)| ≥ 1 were considered significantly differentially expressed. Gene Ontology (GO) enrichment analysis of the differentially expressed genes was performed using GSEAPY (v1.1.3).

### Statistical Analysis

All statistical analyses were performed using GraphPad Prism v9.0 (GraphPad Software). For comparisons between two independent groups, unpaired two-tailed Student’s t-tests were applied when data met the assumptions of normality and homogeneity of variance. For non-parametric data, the Mann–Whitney U test was used. Comparisons involving more than two groups were analyzed using one-way ANOVA followed by appropriate post hoc tests. Data are presented as mean ± SEM (or mean ± SD where indicated), and statistical significance was defined as P < 0.05. Detailed statistical information, including sample sizes (n) and specific tests applied are provided in the corresponding figure legends.

## ACKNOWLEDGMENTS

We thank Dr. Zhiqiang Yan (Chinese Institutes for Medical Research), Dr. Yang Xu (Southern University of Science and Technology) and Dr. Jianzhi Zeng (Shenzhen Bay Laboratory) for generously providing reagents and offering valuable discussions on this project. We thank Xiaojia Wang for assistance on the project. We also thank the assistance of SUSTech Core Research Facilities on flow cytometry and imaging.

## AUTHOR CONTRIBUTIONS

R.T. conceived and designed the study. M.O. and J.W. performed all experiments and data analyses under the guidance of X.L. and R.T. M.O. and R.T. wrote the manuscript with input from all authors.

## SUPPLEMENTAL INFORMATION

Document S1: Figures S1–S5 and their legends

Table S1: Raw sgRNA counts of the CaRPOOL screen

Table S2: MAGeCK-iNC results of the CaRPOOL screen

Table S3: Primer and plasmid sequences used in this study.

## References

1. Galluzzi L, Yamazaki T, Kroemer G: Linking cellular stress responses to systemic homeostasis. Nat Rev Mol Cell Biol 2018, 19:731–745.

2. Ochoa CD, Wu RF, Terada LS: ROS signaling and ER stress in cardiovascular disease. Mol Aspects Med 2018, 63:18–29.

3. Zhang W, Wu Y, S JG: Membrane adhesion junctions regulate airway smooth muscle phenotype and function. Physiol Rev 2023, 103:2321–2347.

4. Kefauver JM, Ward AB, Patapoutian A: Discoveries in structure and physiology of mechanically activated ion channels. Nature 2020, 587:567–576.

5. Scholz N, Dahse AK, Kemkemer M, Bormann A, Auger GM, Vieira Contreras F, Ernst LF, Staake H, Korner MB, Buhlan M, et al: Molecular sensing of mechano- and ligand-dependent adhesion GPCR dissociation. Nature 2023, 615:945–953.

6. Caterina MJ, Schumacher MA, Tominaga M, Rosen TA, Levine JD, Julius D: The capsaicin receptor: a heat-activated ion channel in the pain pathway. Nature 1997, 389:816–824.

7. Vriens J, Nilius B, Voets T: Peripheral thermosensation in mammals. Nat Rev Neurosci 2014, 15:573–589.

8. Shi Y, Hu J, Xue T: Light, opsins, and life: Mammalian photophysiological functions beyond image perception. Neuron 2025, 113:3108–3128.

9. Billesbolle CB, de March CA, van der Velden WJC, Ma N, Tewari J, Del Torrent CL, Li L, Faust B, Vaidehi N, Matsunami H, Manglik A: Structural basis of odorant recognition by a human odorant receptor. Nature 2023, 615:742–749.

10. Delventhal R, Carlson JR: Bitter taste receptors confer diverse functions to neurons. Elife 2016, 5.

11. Ranade SS, Syeda R, Patapoutian A: Mechanically Activated Ion Channels. Neuron 2015, 87:1162–1179.

12. Coste B, Mathur J, Schmidt M, Earley TJ, Ranade S, Petrus MJ, Dubin AE, Patapoutian A: Piezo1 and Piezo2 are essential components of distinct mechanically activated cation channels. Science 2010, 330:55–60.

13. Gong J, Liu J, Ronan EA, He F, Cai W, Fatima M, Zhang W, Lee H, Li Z, Kim GH, et al: A Cold-Sensing Receptor Encoded by a Glutamate Receptor Gene. Cell 2019, 178:1375–1386 e1311.

14. Xu J, Mathur J, Vessieres E, Hammack S, Nonomura K, Favre J, Grimaud L, Petrus M, Francisco A, Li J, et al: GPR68 Senses Flow and Is Essential for Vascular Physiology. Cell 2018, 173:762–775 e716.

15. Leng K, Kampmann M: Towards elucidating disease-relevant states of neurons and glia by CRISPR-based functional genomics. Genome Med 2022, 14:130.

16. Das S, Bano S, Kapse P, Kundu GC: CRISPR based therapeutics: a new paradigm in cancer precision medicine. Mol Cancer 2022, 21:85.

17. Vercauteren S, Fiesack S, Maroc L, Verstraeten N, Dewachter L, Michiels J, Vonesch SC: The rise and future of CRISPR-based approaches for high-throughput genomics. FEMS Microbiol Rev 2024, 48.

18. Li K, Ouyang M, Zhan J, Tian R: CRISPR-based functional genomics screening in human-pluripotent-stem-cell-derived cell types. Cell Genom 2023, 3:100300.

19. de Bakker V, Liu X, Bravo AM, Veening JW: CRISPRi-seq for genome-wide fitness quantification in bacteria. Nat Protoc 2022, 17:252–281.

20. Franks SNJ, Heon-Roberts R, Ryan BJ: CRISPRi: a way to integrate iPSC-derived neuronal models. Biochem Soc Trans 2024, 52:539–551.

21. Zhang Y, Rozsa M, Liang Y, Bushey D, Wei Z, Zheng J, Reep D, Broussard GJ, Tsang A, Tsegaye G, et al: Fast and sensitive GCaMP calcium indicators for imaging neural populations. Nature 2023, 615:884–891.

22. Fosque BF, Sun Y, Dana H, Yang CT, Ohyama T, Tadross MR, Patel R, Zlatic M, Kim DS, Ahrens MB, et al: Neural circuits. Labeling of active neural circuits in vivo with designed calcium integrators. Science 2015, 347:755–760.

23. Moeyaert B, Holt G, Madangopal R, Perez-Alvarez A, Fearey BC, Trojanowski NF, Ledderose J, Zolnik TA, Das A, Patel D, et al: Improved methods for marking active neuron populations. Nat Commun 2018, 9:4440.

24. Yuan F, Yang H, Xue Y, Kong D, Ye R, Li C, Zhang J, Theprungsirikul L, Shrift T, Krichilsky B, et al: OSCA1 mediates osmotic-stress-evoked Ca2+ increases vital for osmosensing in Arabidopsis. Nature 2014, 514:367–371.

25. Centeio R, Ousingsawat J, Schreiber R, Kunzelmann K: Ca(2+) Dependence of Volume-Regulated VRAC/LRRC8 and TMEM16A Cl(-) Channels. Front Cell Dev Biol 2020, 8:596879.

26. Inoue R, Jensen LJ, Jian Z, Shi J, Hai L, Lurie AI, Henriksen FH, Salomonsson M, Morita H, Kawarabayashi Y, et al: Synergistic activation of vascular TRPC6 channel by receptor and mechanical stimulation via phospholipase C/diacylglycerol and phospholipase A2/omega-hydroxylase/20-HETE pathways. Circ Res 2009, 104:1399–1409.

27. Zhang M, Wang D, Kang Y, Wu JX, Yao F, Pan C, Yan Z, Song C, Chen L: Structure of the mechanosensitive OSCA channels. Nat Struct Mol Biol 2018, 25:850–858.

28. Liu X, Ong HL, Pani B, Johnson K, Swaim WB, Singh B, Ambudkar I: Effect of cell swelling on ER/PM junctional interactions and channel assembly involved in SOCE. Cell Calcium 2010, 47:491–499.

29. Wei T, Li J, Lei X, Lin R, Wu Q, Zhang Z, Shuai S, Tian R: Multimodal CRISPR screens uncover DDX39B as a global repressor of A-to-I RNA editing. Cell Rep 2025, 44:116009.

30. Horlbeck MA, Gilbert LA, Villalta JE, Adamson B, Pak RA, Chen Y, Fields AP, Park CY, Corn JE, Kampmann M, Weissman JS: Compact and highly active next-generation libraries for CRISPR-mediated gene repression and activation. Elife 2016, 5.

31. Jeong H, Clark S, Goehring A, Dehghani-Ghahnaviyeh S, Rasouli A, Tajkhorshid E, Gouaux E: Structures of the TMC-1 complex illuminate mechanosensory transduction. Nature 2022, 610:796–803.

32. Fu S, Pan X, Lu M, Dong J, Yan Z: Human TMC1 and TMC2 are mechanically gated ion channels. Neuron 2025, 113:411–425 e414.

33. Guan Y, Du HB, Yang Z, Wang YZ, Ren R, Liu WW, Zhang C, Zhang JH, An WT, Li NN, et al: Deafness-Associated ADGRV1 Mutation Impairs USH2A Stability through Improper Phosphorylation of WHRN and WDSUB1 Recruitment. Adv Sci (Weinh) 2023, 10:e2205993.

34. Ligezowska A, Boye K, Eble JA, Hoffmann B, Klosgen B, Merkel R: Mechanically enforced bond dissociation reports synergistic influence of Mn2+ and Mg2+ on the interaction between integrin alpha7beta1 and invasin. J Mol Recognit 2011, 24:715–723.

35. de Rezende FF, Martins Lima A, Niland S, Wittig I, Heide H, Schroder K, Eble JA: Integrin alpha7beta1 is a redox-regulated target of hydrogen peroxide in vascular smooth muscle cell adhesion. Free Radic Biol Med 2012, 53:521–531.

36. Bao X, Kobayashi M, Hatakeyama S, Angata K, Gullberg D, Nakayama J, Fukuda MN, Fukuda M: Tumor suppressor function of laminin-binding alpha-dystroglycan requires a distinct beta3-N-acetylglucosaminyltransferase. Proc Natl Acad Sci U S A 2009, 106:12109–12114.

37. Hamann J, Aust G, Arac D, Engel FB, Formstone C, Fredriksson R, Hall RA, Harty BL, Kirchhoff C, Knapp B, et al: International Union of Basic and Clinical Pharmacology. XCIV. Adhesion G protein-coupled receptors. Pharmacol Rev 2015, 67:338–367.

38. Lv Z, Ding Y, Cao W, Wang S, Gao K: Role of RHO family interacting cell polarization regulators (RIPORs) in health and disease: Recent advances and prospects. Int J Biol Sci 2022, 18:800–808.

39. Asmar A, Barrett-Jolley R, Werner A, Kelly R, Jr., Stacey M: Membrane channel gene expression in human costal and articular chondrocytes. Organogenesis 2016, 12:94–107.

40. Lukashkina VA, Levic S, Lukashkin AN, Strenzke N, Russell IJ: A connexin30 mutation rescues hearing and reveals roles for gap junctions in cochlear amplification and micromechanics. Nat Commun 2017, 8:14530.

41. Forster R, Davalos-Misslitz AC, Rot A: CCR7 and its ligands: balancing immunity and tolerance. Nat Rev Immunol 2008, 8:362–371.

42. Topaz O, Shurman DL, Bergman R, Indelman M, Ratajczak P, Mizrachi M, Khamaysi Z, Behar D, Petronius D, Friedman V, et al: Mutations in GALNT3, encoding a protein involved in O-linked glycosylation, cause familial tumoral calcinosis. Nat Genet 2004, 36:579–581.

43. Seko A, Hara-Kuge S, Yamashita K: Molecular cloning and characterization of a novel human galactose 3-O-sulfotransferase that transfers sulfate to gal beta 1-->3galNAc residue in O-glycans. J Biol Chem 2001, 276:25697–25704.

44. Liang C, Zhang Q, Chen X, Liu J, Tanaka M, Wang S, Lepler SE, Jin Z, Siemann DW, Zeng B, Tang X: Human cancer cells generate spontaneous calcium transients and intercellular waves that modulate tumor growth. Biomaterials 2022, 290:121823.

45. Albarran-Juarez J, Iring A, Wang S, Joseph S, Grimm M, Strilic B, Wettschureck N, Althoff TF, Offermanns S: Piezo1 and G(q)/G(11) promote endothelial inflammation depending on flow pattern and integrin activation. J Exp Med 2018, 215:2655–2672.

46. Zhao C, MacKinnon R: Structural and functional analyses of a GPCR-inhibited ion channel TRPM3. Neuron 2023, 111:81–91 e87.

47. Choi D, Park E, Yu RP, Cooper MN, Cho IT, Choi J, Yu J, Zhao L, Yum JI, Yu JS, et al: Piezo1-Regulated Mechanotransduction Controls Flow-Activated Lymphatic Expansion. Circ Res 2022, 131:e2–e21.

48. Atcha H, Jairaman A, Holt JR, Meli VS, Nagalla RR, Veerasubramanian PK, Brumm KT, Lim HE, Othy S, Cahalan MD, et al: Mechanically activated ion channel Piezo1 modulates macrophage polarization and stiffness sensing. Nat Commun 2021, 12:3256.

49. Jiang F, Yin K, Wu K, Zhang M, Wang S, Cheng H, Zhou Z, Xiao B: The mechanosensitive Piezo1 channel mediates heart mechano-chemo transduction. Nat Commun 2021, 12:869.

50. Solis AG, Bielecki P, Steach HR, Sharma L, Harman CCD, Yun S, de Zoete MR, Warnock JN, To SDF, York AG, et al: Mechanosensation of cyclical force by PIEZO1 is essential for innate immunity. Nature 2019, 573:69–74.

51. Baratchi S, Danish H, Chheang C, Zhou Y, Huang A, Lai A, Khanmohammadi M, Quinn KM, Khoshmanesh K, Peter K: Piezo1 expression in neutrophils regulates shear-induced NETosis. Nat Commun 2024, 15:7023.

52. Zhang T, Bi C, Li Y, Zhao L, Cui Y, Ouyang K, Xiao B: Phosphorylation of Piezo1 at a single residue, serine-1612, regulates its mechanosensitivity and in vivo mechanotransduction function. Neuron 2024, 112:3618–3633 e3616.

53. Iring A, Jin YJ, Albarran-Juarez J, Siragusa M, Wang S, Dancs PT, Nakayama A, Tonack S, Chen M, Kunne C, et al: Shear stress-induced endothelial adrenomedullin signaling regulates vascular tone and blood pressure. J Clin Invest 2019, 129:2775–2791.

54. Liu W, Liu C, Ren PG, Chu J, Wang L: An Improved Genetically Encoded Fluorescent cAMP Indicator for Sensitive cAMP Imaging and Fast Drug Screening. Front Pharmacol 2022, 13:902290.

55. Wang L, Wu C, Peng W, Zhou Z, Zeng J, Li X, Yang Y, Yu S, Zou Y, Huang M, et al: A high-performance genetically encoded fluorescent indicator for in vivo cAMP imaging. Nat Commun 2022, 13:5363.

56. Alraies Z, Rivera CA, Delgado MG, Sanseau D, Maurin M, Amadio R, Maria Piperno G, Dunsmore G, Yatim A, Lacerda Mariano L, et al: Cell shape sensing licenses dendritic cells for homeostatic migration to lymph nodes. Nat Immunol 2024, 25:1193–1206.

57. Clatworthy MR, Aronin CE, Mathews RJ, Morgan NY, Smith KG, Germain RN: Immune complexes stimulate CCR7-dependent dendritic cell migration to lymph nodes. Nat Med 2014, 20:1458–1463.

58. Liu J, Zhang X, Chen K, Cheng Y, Liu S, Xia M, Chen Y, Zhu H, Li Z, Cao X: CCR7 Chemokine Receptor-Inducible lnc-Dpf3 Restrains Dendritic Cell Migration by Inhibiting HIF-1alpha-Mediated Glycolysis. Immunity 2019, 50:600–615 e615.

59. Alanko J, Ucar MC, Canigova N, Stopp J, Schwarz J, Merrin J, Hannezo E, Sixt M: CCR7 acts as both a sensor and a sink for CCL19 to coordinate collective leukocyte migration. Sci Immunol 2023, 8:eadc9584.

60. Huo Y, Zhou Y, Lin L, Yang F, Meng J, He F, Zhang F, Song M, Shen C, Liu Y, et al: GABA-independent activation of GABA(B) receptor by mechanical forces. Nat Commun 2025, 16:9843.

61. Fu C, Huang W, Tang Q, Niu M, Guo S, Langenhan T, Song G, Yan J: Unveiling Mechanical Activation: GAIN Domain Unfolding and Dissociation in Adhesion GPCRs. Nano Lett 2023, 23:9179–9186.

62. Mitgau J, Franke J, Schinner C, Stephan G, Berndt S, Placantonakis DG, Kalwa H, Spindler V, Wilde C, Liebscher I: The N Terminus of Adhesion G Protein-Coupled Receptor GPR126/ADGRG6 as Allosteric Force Integrator. Front Cell Dev Biol 2022, 10:873278.

63. Petersen SC, Luo R, Liebscher I, Giera S, Jeong SJ, Mogha A, Ghidinelli M, Feltri ML, Schoneberg T, Piao X, Monk KR: The adhesion GPCR GPR126 has distinct, domain-dependent functions in Schwann cell development mediated by interaction with laminin-211. Neuron 2015, 85:755–769.

64. Huse M: Mechanical forces in the immune system. Nat Rev Immunol 2017, 17:679–690.

65. Angeli V, Lim HY: Biomechanical control of lymphatic vessel physiology and functions. Cell Mol Immunol 2023, 20:1051–1062.

66. Liu J, Zhang X, Cheng Y, Cao X: Dendritic cell migration in inflammation and immunity. Cell Mol Immunol 2021, 18:2461–2471.

67. Kiermaier E, Moussion C, Veldkamp CT, Gerardy-Schahn R, de Vries I, Williams LG, Chaffee GR, Phillips AJ, Freiberger F, Imre R, et al: Polysialylation controls dendritic cell trafficking by regulating chemokine recognition. Science 2016, 351:186–190.

68. Niu L, Cheng B, Huang G, Nan K, Han S, Ren H, Liu N, Li Y, Genin GM, Xu F: A positive mechanobiological feedback loop controls bistable switching of cardiac fibroblast phenotype. Cell Discov 2022, 8:84.

69. Wu J, Chen Y, Liao Z, Liu H, Zhang S, Zhong D, Qiu X, Chen T, Su D, Ke X, et al: Self-amplifying loop of NF-kappaB and periostin initiated by PIEZO1 accelerates mechano-induced senescence of nucleus pulposus cells and intervertebral disc degeneration. Mol Ther 2022, 30:3241–3256.

70. Carreras-Sureda A, Zhang X, Laubry L, Brunetti J, Koenig S, Wang X, Castelbou C, Hetz C, Liu Y, Frieden M, Demaurex N: The ER stress sensor IRE1 interacts with STIM1 to promote store-operated calcium entry, T cell activation, and muscular differentiation. Cell Rep 2023, 42:113540.

71. Kouba S, Zhang X, Nere R, Castelbou C, Demaurex N, Carreras-Sureda A: Campari2 genomic interrogation of homeostatic calcium activity identifies TIM1 as a negative regulator of T cell function. Cell Calcium 2025, 129:103036.

72. Boggess SC, Gandhi V, Tsai MC, Marzette E, Teyssier N, Yu-Ying Chou J, Hu X, Cramer A, Yadanar L, Shroff K, et al: A Massively Parallel CRISPR-Based Screening Platform for Modifiers of Neuronal Activity. bioRxiv 2025.

73. Tian R, Gachechiladze MA, Ludwig CH, Laurie MT, Hong JY, Nathaniel D, Prabhu AV, Fernandopulle MS, Patel R, Abshari M, et al: CRISPR Interference-Based Platform for Multimodal Genetic Screens in Human iPSC-Derived Neurons. Neuron 2019, 104:239–255 e212.

74. Tian R, Abarientos A, Hong J, Hashemi SH, Yan R, Drager N, Leng K, Nalls MA, Singleton AB, Xu K, et al: Genome-wide CRISPRi/a screens in human neurons link lysosomal failure to ferroptosis. Nat Neurosci 2021, 24:1020–1034.

