## Supplementary material for "CaRPOOL: A Pooled Calcium-Recording CRISPR Screening Platform Identifies CCR7 as a Modulator of Cellular Osmomechanosensing": Document S1

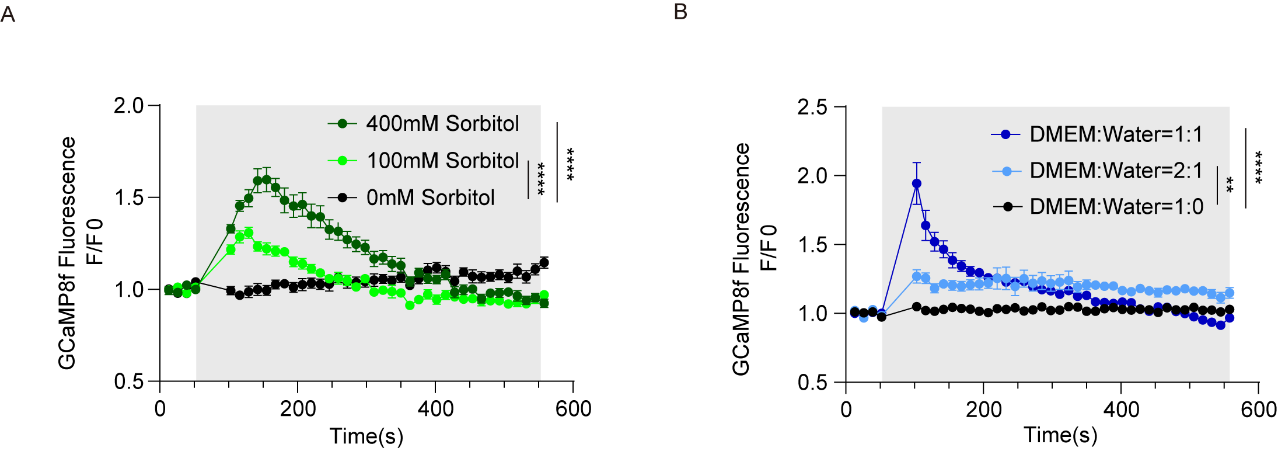

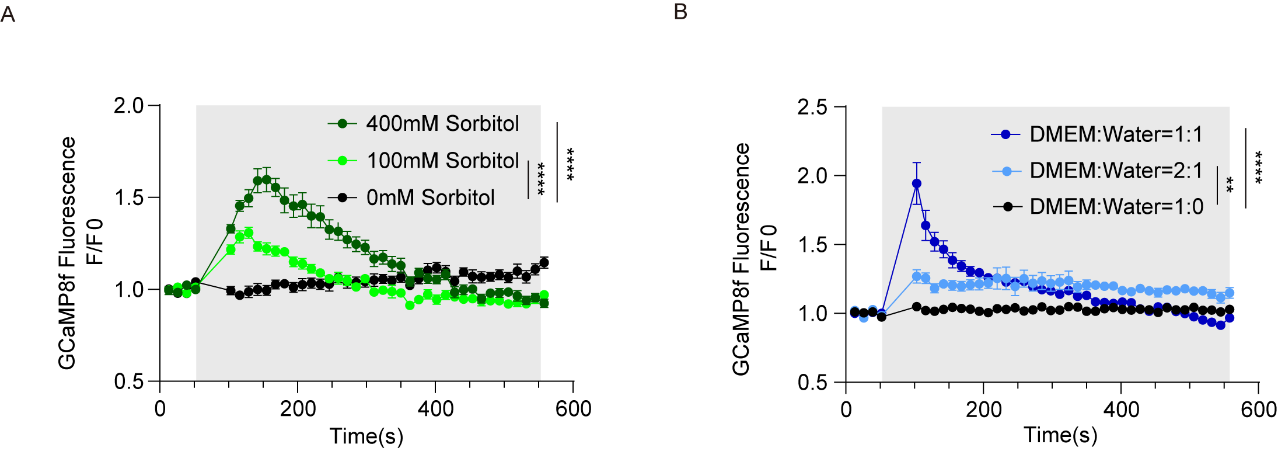
**Figure S1. Calcium responses of HEK293T cells to hypotonic and hypertonic treatments**

(A-B) Calcium responses of HEK293T cells to (A) hypertonic (100 mM and 400 mM sorbitol in DMEM) or (B) hypotonic (2:1 and 1:1 DMEM:ddH₂O, ~200mOsm and ~150mOsm) stimulation monitored by microplate reader using GCaMP8f. Traces show mean F/F₀ ± SEM (n = 10 biological replicates). Gray shading indicates stimulation. **p < 0.01, ***p < 0.001, ****p < 0.0001. One-way ANOVA.

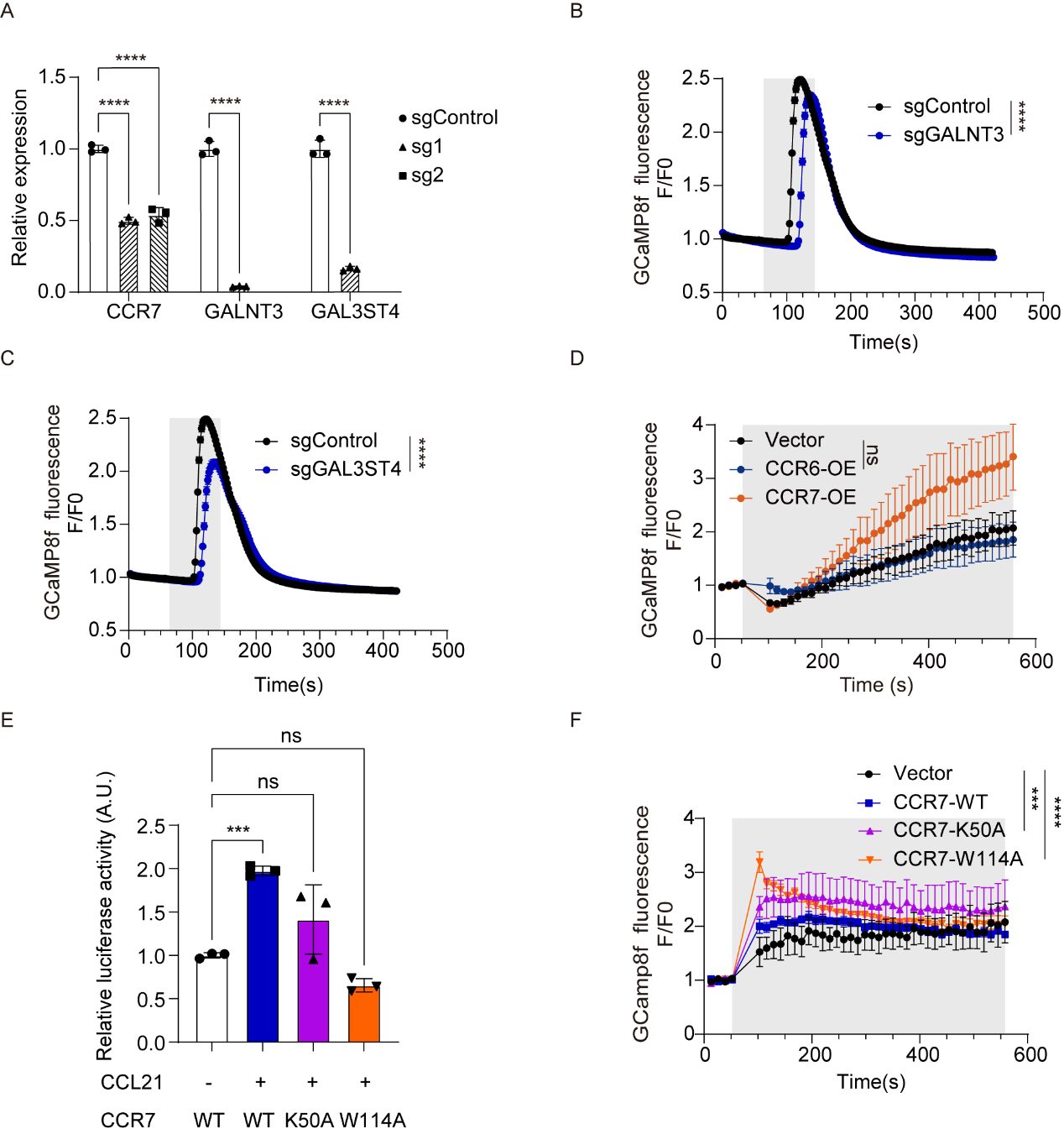

**Figure S2. Validation of top hits from the CaMPARI2-based CRISPRi screen reveals *CCR7* as a key regulator of osmomechanical signaling**

(A) Validation of CRISPRi-mediated gene knockdown. Relative mRNA expression levels of *CCR7*, *GALNT3*, and *GAL3ST4* in HEK293T cells expressing sgControl or gene-targeting sgRNAs were measured by qRT-PCR. Data are mean ± SEM (n = 3 technical replicates). ****p < 0.0001. Unpaired two‑tailed Student’s t-test.

(B-C) Knockdown of *GAL3ST4* or *GALNT3* reduces calcium responses to hypotonic stimulation. GCaMP8f fluorescence traces of HEK293T cells expressing sgControl or sgGAL3ST4 (left) and sgControl or sgGALNT3 (right), measured by fluorescence imaging. Traces show mean F/F₀ ± SEM (n = 297 cells for sgControl, n = 196 cells for sgGAL3ST4, n = 312 cells sgGALNT3). Gray shading indicates stimulation. ****p < 0.0001. Unpaired two‑tailed Student’s t-test.

(D) CCR7, but not CCR6, enhances osmomechanical calcium signaling. GCaMP8f fluorescence traces of HEK293T cells expressing empty vector, *CCR6*, or *CCR7* were measured by microplate reader. Traces show mean F/F₀ ± SEM (n = 5 biological replicates). Gray shading indicates stimulation. ns, not significant. Unpaired two‑tailed Student’s t-test.

(E) Relative luciferase activity indicating CCR7 signaling in response to CCL21 stimulation. Cells expressing wild-type CCR7 (CCR7-WT) show increased activity with CCL21, while CCR7 mutants (CCR7-K50A, CCR7-W114A) that abolish CCL21 binding do not.Data are expressed in RLU (relative luminescence units) for the vector, *CCR7*, *CCR7-K50A* or *CCR7-W114A* (n = 3 biological replicate). ns, not significant, ***p < 0.001. One-way ANOVA.

(F) GCaMP8f fluorescence (F/F0) traces illustrating intracellular calcium responses over time in cells expressing vector, wild-type CCR7, or the CCR7-K50A and CCR7-W114A mutants under osmomechanical stimulation (shaded area) measured by microplate reader. Traces represent mean F/F₀ ± SEM (n = 4 biological replicates). ***p < 0.001, ****p < 0.0001. One‑way ANOVA.

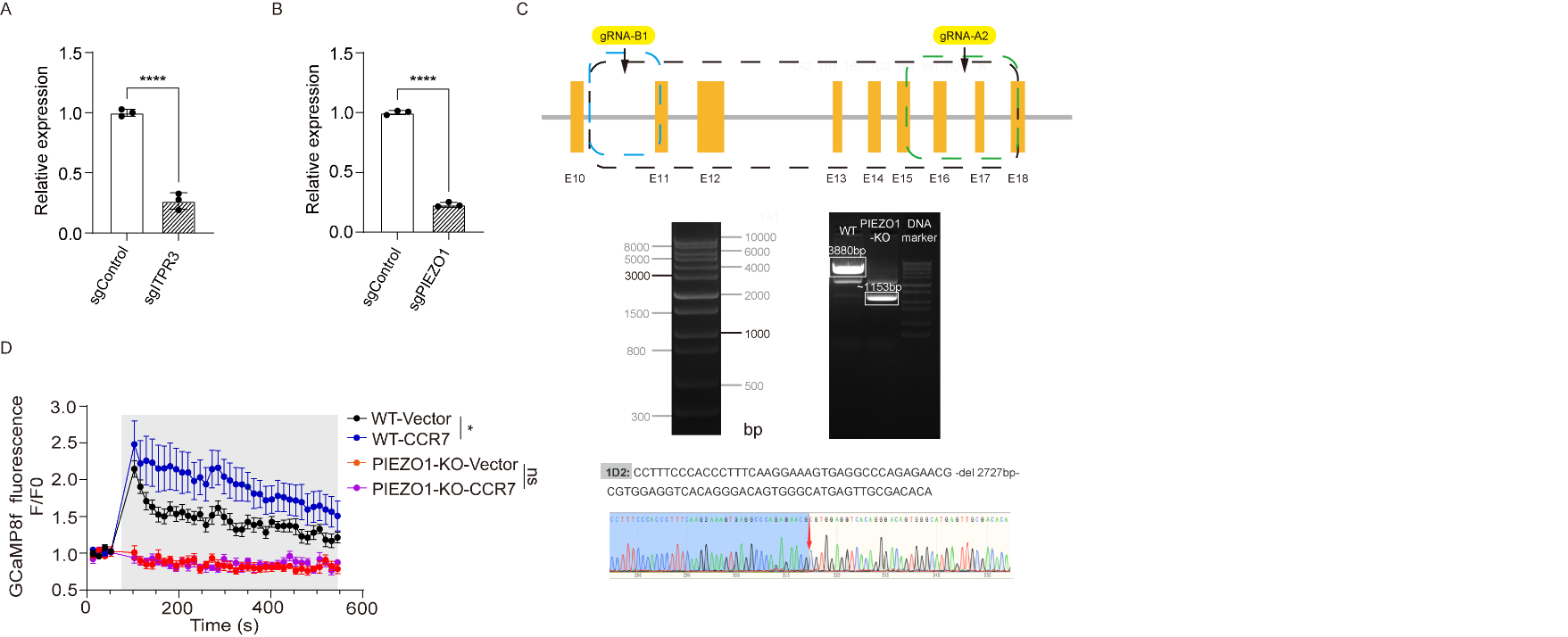

**Figure S3. CCR7 regulates osmomechanical calcium signaling through PIEZO1**

(A–B) Validation of CRISPRi-mediated knockdown efficiency. Relative mRNA expression of *ITPR3* (A) and *PIEZO1* (B) in HEK293T cells expressing sgControl or gene-specific sgRNAs, measured by qRT-PCR. Data are mean ± SD (n = 3 technical replicates). ****p < 0.0001. Unpaired two‑tailed Student's t-test.

(C) PCR and Sanger sequencing validation of *PIEZO1*-KO cells.

(D) *CCR7* overexpression fails to restore calcium influx in *PIEZO1*‑knockout cells. GCaMP8f traces of WT or PIEZO1-KO cells transfected with vector or *CCR7* are shown alongside sgControl‑Vector cells, measured by microplate reader. Traces represent mean F/F₀ ± SEM (n = 5 biological replicates). Gray shading indicates stimulation. ns, not significant, *p < 0.05. One‑way ANOVA.

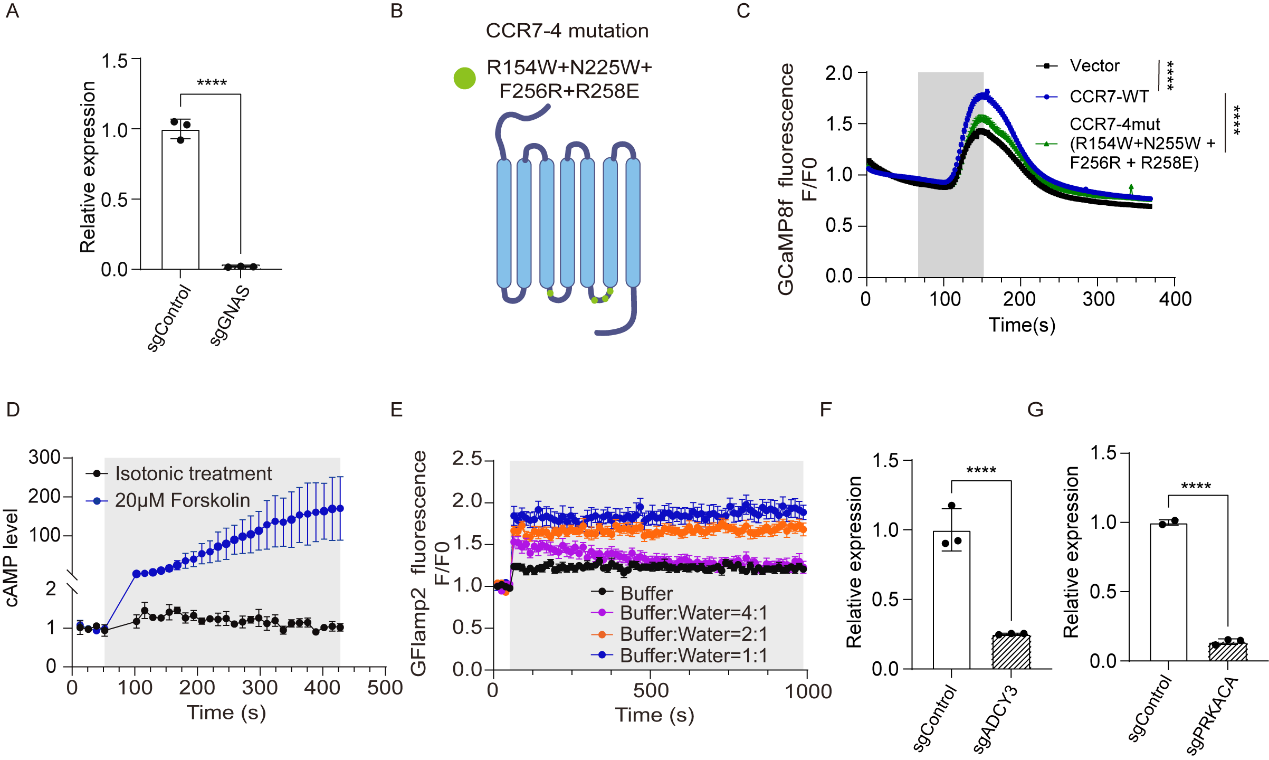

**Figure S4. CCR7 modulates osmomechanical calcium signaling through the Gαs-cAMP-PKA Pathway**

(A) Validation of CRISPRi-mediated knockdown efficiency. Relative mRNA expression of *GNAS* in HEK293T cells expressing sgControl or *GNAS*-targeting sgRNA (sgGNAS), measured by qRT-PCR. Data are mean ± SD (n = 3 technical replicates). ****p < 0.0001. Unpaired two‑tailed Student's t-test.

(B) Schematic representation of mutations in the CCR7.

(C) GCaMP8f fluorescence (F/F0) traces illustrating intracellular calcium responses over time in cells expressing vector, wild-type CCR7, or the CCR7-4 mutations under osmomechanical stimulation (shaded area) measured by fluorescence imaging. Traces represent mean F/F₀ ± SEM (n = 4 biological replicates). ****p < 0.0001. One‑way ANOVA.

(D) Forskolin activates adenylyl cyclase and increases intracellular cAMP levels. HEK293T cells expressing the cAMP biosensor GloSensor-22F were treated with isotonic buffer or 20 μM forskolin. Traces show mean cAMP levels ± SEM (n = 2 or 3 biological replicates), measured by microplate reader. Gray shading indicates treatment period.

(E) Hypotonic stimulation increases intracellular cAMP levels in an osmolarity-dependent manner. HEK293T cells expressing the genetically encoded cAMP sensor GFlamp2 were subjected to varying degrees of hypotonic stimulation (buffer:water ratios of 4:1, 2:1, and 1:1). Traces show mean fluorescence intensity (F/F₀) ± SEM (n = 10 biological replicates), measured by microplate reader. Gray shading indicates stimulation period.

(F-G) Validation of CRISPRi-mediated knockdown efficiency. Relative mRNA expression of *ADCY3* (F) and *PRKACA* (G) in HEK293T cells expressing sgControl or gene-specific sgRNAs, measured by qRT-PCR. Data are mean ± SD (n = 3 technical replicates). Unpaired two‑tailed Student's t-test.

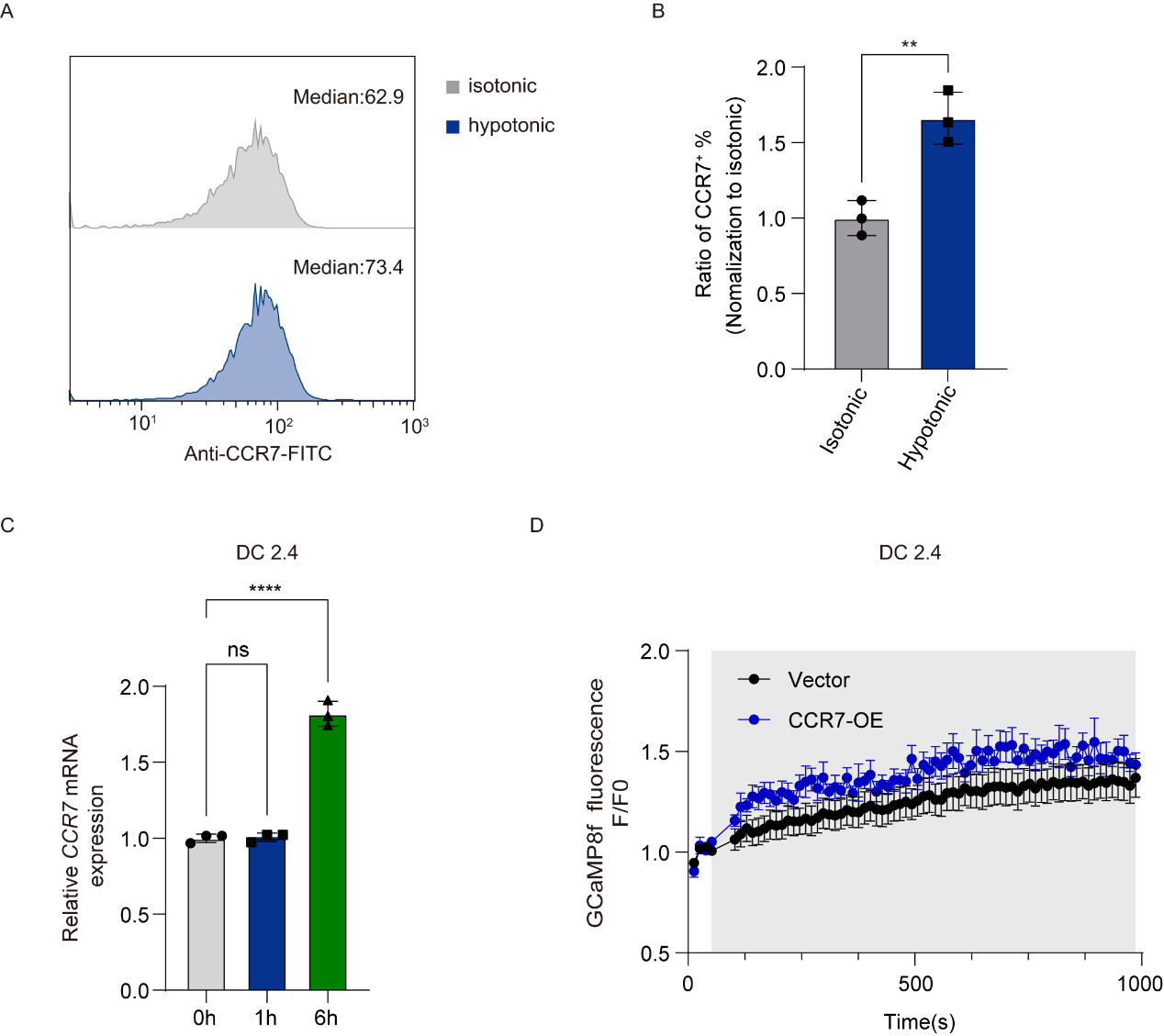

**Figure S5. CCR7 Expression is Induced by Osmomechanical Stimulation and Enhances Calcium Signaling in Immune Cell Lines**

(A) Flow cytometry histograms of CCR7-FITC surface staining in Jurkat cells under isotonic versus hypotonic treatment, showing increased CCR7 surface expression after hypotonic exposure (two-color overlay with median indicated).

(B) Quantification of surface CCR7 expression as the ratio of hypotonic to isotonic MFI (Jurkat cells). Data are mean ± SD (n = 3 biological replicates). **p < 0.01. Unpaired two‑tailed Student's t-test.

(C) Hypotonic stimulation upregulates *CCR7* mRNA expression in DC2.4 cells. Relative *CCR7* mRNA levels at 0, 4, and 6 hours after hypotonic treatment, measured by qRT-PCR. Data are mean ± SD (n = 3 technical replicates). ns, not significant, ****p < 0.0001. One-way ANOVA.

(D) *CCR7* overexpression enhances calcium responses to hypotonic stimulation in DC2.4 cells. Fluo-4 fluorescence traces of cells transfected with empty vector or *CCR7*, measured by microplate reader. Traces show mean F/F₀ ± SEM (n = 8 biological replicates). Gray shading indicates stimulation.
